# Early-life experience reorganizes neuromodulatory regulation of stage-specific behavioral responses and individuality types during development

**DOI:** 10.1101/2022.10.24.513603

**Authors:** Reemy Ali Nasser, Yuval Harel, Shay Stern

## Abstract

Early-life experiences may promote stereotyped behavioral alterations that are dynamic across development time, but also behavioral responses that are variable among individuals, even when initially exposed to the same stimulus. Here, by utilizing longitudinal monitoring of *C. elegans* individuals throughout development we show that behavioral effects of early-life starvation are exposed during early and late developmental stages and buffered during intermediate stages of development. We further found that both dopamine and serotonin shape the discontinuous behavioral responses by opposite and temporally segregated functions across development time. While dopamine buffers behavioral responses during intermediate developmental stages, serotonin promotes behavioral sensitivity to stress during early and late stages. Interestingly, unsupervised analysis of individual biases across development uncovered multiple individuality types that coexist within stressed and unstressed populations and further identified experience-dependent effects on their composition. These results provide insight into the complex temporal regulation of behavioral plasticity across developmental timescales, structuring shared and unique individual responses to early-life experiences.

## Introduction

Long-term behavioral patterns across development are highly dynamic across and within developmental stages and are temporally synchronized with the individual’s developmental clock. For instance, flies show differences in foraging behavior that depend on their larval stage (Sokolowski et al., 1984), fish internally modify their startle response across life (Kimmel et al., 1974) and stomatogastric motor patterns are regulated across development (Rehm et al., 2008). In addition, fear-extinction learning is inhibited during adolescence compared to other life-stages in humans and mice (Pattwell et al., 2012). Long-term behavioral outputs are influenced by the changes in the internal state of individuals, as well as by their past and current environmental exposures. In particular, animals may be transiently exposed to environmental perturbations at different stages of life, but experiences during early developmental windows, which are usually referred to as critical or sensitive periods, were shown to generate long-lasting effects (Lorenz, 1935; Korosi et al., 2012; Jin et al., 2016; Nevitt et al., 1994; Remy and Hobert, 2005). This stable imprinting of early memories has the potential to increase survival and reproduction during later life-stages of the organism (Immelmann, 1975). However, a complete temporal view of the long-term effects of early experiences on behavior throughout development, across and within all stages, is still lacking.

While long-lasting effects on behavior may be shared by many individuals, reflected by stereotypic behavioral responses following an early-life experience, individuals within the same population may also show unique patterns of long-term behavior that distinguish them from each other. This inter-individual variation in behavioral responses may be exposed even when animals are initially experiencing the same early conditions. Here we study how early-life experiences shape stage-specific behavioral patterns across development and how they affect the diversity in long-term behavioral responses among individuals. Consistent behavioral individuality within isogenic populations that were raised in the same environment has been previously described in various species, including in the pea aphid (Schuett et al., 2011), *D. melanogaster* (Buchanan et al., 2015; Kain et al., 2012; Linneweber et al., 2020), clonal fish (Bierbach et al., 2017) and mice (Freund et al., 2013). The nematode *C. elegans* in an ideal system to study how early-life experiences shape long-term behavior and inter-individual variation across developmental timescales due to their short development time of 2.5 days and the homogeneous populations generated by the self-fertilizing reproduction mode of the hermaphrodite. It was previously shown that under normal growth conditions, *C. elegans* shows both stereotypic patterns of long-term behavior and consistent individual biases within the isogenic population (Stern et al., 2017).

By continuously tracking the locomotory behavior of single individuals following transient periods of starvation early in life throughout their complete developmental trajectory, we show that early-life starvation exposes long-term behavioral plasticity that is discontinuous over development time. Temporal differences in long-term behavioral responses to early stress are reflected by strong behavioral modifications during early and late developmental stages and the buffering of behavioral effects during intermediate stages of development. We further found that dopamine maintains the buffering of behavioral responses during mid-development and serotonin promotes behavioral sensitivity to early starvation during early and late stages of development. Moreover, by performing unsupervised analysis of patterns of individual biases across development, we identified a spectrum of temporal individuality types that are dominant within stressed and unstressed populations. Both the early-life history and neuromodulatory state of the population affect variation in specific individuality types. These results show how a transient early-life environment shapes a long-term behavioral structure of stereotypic and variable responses across developmental stages.

## Results

### Early-life stress generates discontinuous and distinct behavioral effects at different stages of development

To study how stressful environments early in life influence long-term behavioral patterns across and within all developmental stages we continuously tracked the behavior of individual animals following a transient period of starvation early in development. Imaging was performed using a custom-made multi-camera imaging system across the full developmental trajectory of *C. elegans* individuals (55 hours), at high spatiotemporal resolution (3fps, ~10um) and in a tightly controlled environment (Stern et al., 2017). First larval stage (L1) animals that hatch into an environment that completely lacks a food source do not grow and their development is arrested (Greenwald and Horvitz, 1982; Johnson et al., 1984; Baugh, 2013). Following L1 arrest, when animals encounter food, they resume their normal developmental trajectory to reach adulthood. This early-stress paradigm allows us to maintain a homogeneous stress environment across individuals at their earliest stage of development, immediately after hatching.

We continuously monitored single N2 wild-type individuals grown in isolation from their first larval stage to 16 hours of adulthood (n=456) on defined concentrations of UV-killed OP50 bacteria, following periods of stress ranging from 1 to 4 days of early starvation (Fig. 1A,B). In parallel, we tracked the behavior of individuals grown continuously on food, without experiencing starvation (Fig. 1A,B). Animals exposed to early starvation required more time to complete their development (Fig. S1A). To align developmental trajectories of different individuals in time, we age-normalized individuals by dividing each developmental stage, detected by the lethargus period during molting (Cassada and Russell, 1975), into 75 time windows (Fig. S1C) (Stern et al., 2017). Whilst growing in a food environment, *C. elegans* shifts between two behavioral states called roaming and dwelling that last seconds to minutes (Arous et al., 2009; Flavell et al., 2013; Fujiwara et al., 2002; Stern et al., 2017). During a roaming episode animals explore a large area by high-speed forward movements, while in the dwelling episode they show dramatically less exploration due to low-speed movements coupled with frequent reorientations (Fig. S1B). We quantified long-term patterns of locomotory behavior shown by individuals throughout development by measuring two behavioral parameters: fraction of time spent roaming, and speed during roaming episodes.

**Figure 1.**
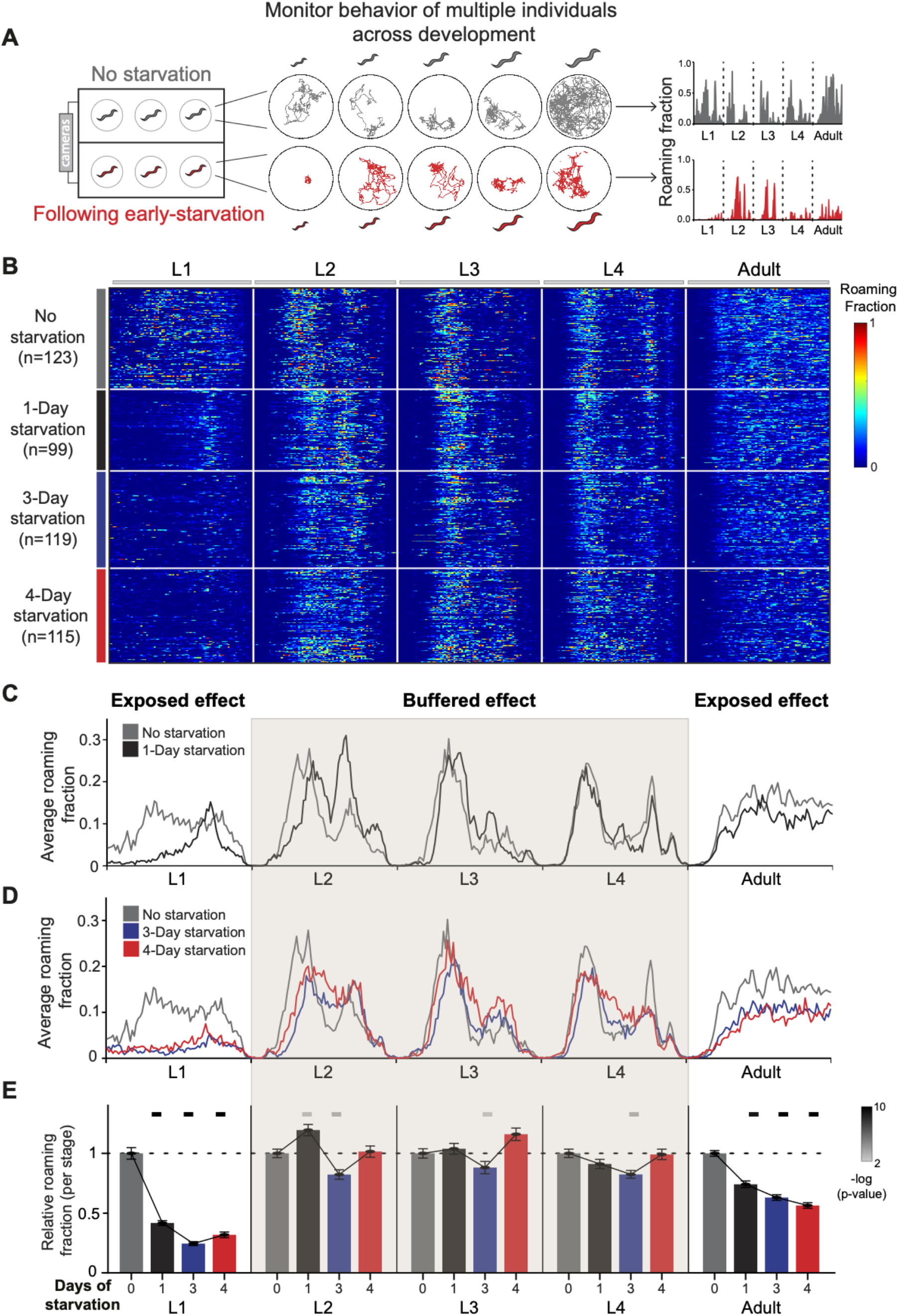
Long-term behavioral tracking of *C. elegans* following early starvation reveals discontinuous behavioral effects across developmental stages. **(A)** Multi-camera imaging system allows longitudinal behavioral tracking of multiple individual worms across all stages of development following early L1 starvation and without starvation, under tightly controlled environmental conditions. Shown are representative locomotion trajectories (middle) and age-normalized roaming activity (right) of post-starved (red) and well-fed (grey) individual worms across all four larval stages and adulthood. Normalization equally divides each stage into 75 time bins. **(B)** Roaming and dwelling behavior of wild-type N2 animals without early starvation (n=123) and following 1-day (n=99), 3-day (n=119) and 4-day starvation (n=115). Each row indicates the age-normalized behavior of one individual across all developmental stages. The different stages are separated by vertical white lines indicating the middle of the lethargus state. Color bar represents the fraction of time spent roaming in each of the 375 time-bins. **(C)** Average roaming fraction of 1-day starved wild-type animals compared to the unstarved population. **(D)** Average roaming fraction of 3- and 4-day starved wild-type animals compared to the unstarved population. **(E)** Average roaming fraction relative to the unstarved population in each developmental stage. Shading highlights intermediate stages of development in which average behavioral effects within a stage are buffered. Error bars indicate standard error of the mean. Upper bars indicate statistical significance (Wilcoxon rank-sum test, FDR corrected) of difference in average roaming fraction between starved and unstarved populations (-log(P-value), indicated are P-values<0.01).

Unstressed individuals hatching in a food environment show dynamic behavioral structures of roaming activity across development, as was previously shown (Stern et al., 2017) (Fig. 1B-D). We found that a transient exposure to early-life starvation generates alterations in long-term behavioral patterns throughout development that were distinct and discontinuous across and within developmental stages. A short early-starvation period of 1 day strongly decreased average roaming activity levels during the L1 and adult stages compared to unstressed individuals (Fig. 1C,E; S1D). In contrast, while 1 day of early starvation modified the temporal structure of activity peaks within the L2 stage, overall roaming activity was not decreased during this stage. Similarly, we found that roaming activity levels were also maintained during the L3 and L4 stages, following 1-day starvation (Fig. 1C,E; S1D). Interestingly, animals exposed to longer starvation periods of 3 and 4 days further showed strong roaming decrease during L1 and adulthood (Figure 1D,E), but exhibited only minor effects on roaming activity within the intermediate L2,L3 and L4 larval stages (Fig. 1D-E; S1D). These results indicate that, while the memory of early starvation is maintained throughout development to expose strong decrease in overall roaming behavior during L1 and adulthood, behavioral effects are buffered across intermediate development times.

Similar to the stage-specific effects of early starvation on the fraction of time spent roaming, instantaneous speed during roaming episodes in individuals exposed to early starvation was affected more strongly at the L1 and adult stages, compared to the intermediate stages (Figure S1E). In summary, transient early-life starvation discontinuously reshapes long-term behavioral patterns across development time by exposing strong behavioral alterations at early and late developmental stages and buffered effects during intermediate stages.

### Unsupervised analysis uncovers multiple individuality types within stressed and unstressed populations

The longitudinal measurements in single animals across development allow us to further quantify long-term inter-individual diversity within stressed and unstressed populations. Following early starvation, different individuals show substantial variation in long-term behavioral responses. For instance, during L1 and adulthood, a fraction of wild-type individuals that were exposed to early stress show 8 to 10-fold decrease in roaming activity relative to the average roaming level of the unstressed population, while other stressed individuals show roaming activity which is indistinguishable from unstressed animals (Fig. S1D). Behavioral individuality is usually defined as a consistent tendency of an individual to show the same behavioral bias relative to the population across long time-periods (Bierbach et al., 2017; Buchanan et al., 2015; Kain et al., 2012; Stern et al., 2017; Schuett et al., 2011). However, individuals may also show alternative patterns of temporal behavioral biases relative to the population that are not random and represent more complex structures of individual biases over time. Here we extend the ‘classic’ analysis of individuality and ask if alternative individuality types coexist within *C. elegans* populations across development.

To analyze long-term individual biases in behavior we first systematically rank individuals based on their roaming activity compared to all other individuals within the same experiment across developmental windows (50 time-bins) (Fig. 2A,B). The rank approach allows us to homogeneously compare between individuals at each developmental window. To take an unsupervised approach for detecting temporal patterns of individual biases that are dominant within stressed and unstressed populations we performed principal component analysis (PCA) of the temporal behavioral ranks of all wild-type individuals (n=456). We then identified statistically significant PC dimensions by comparing them to PCs obtained from a randomly shuffled rank dataset (Fig. 2C-F; Fig. S2A).

**Figure 2.**
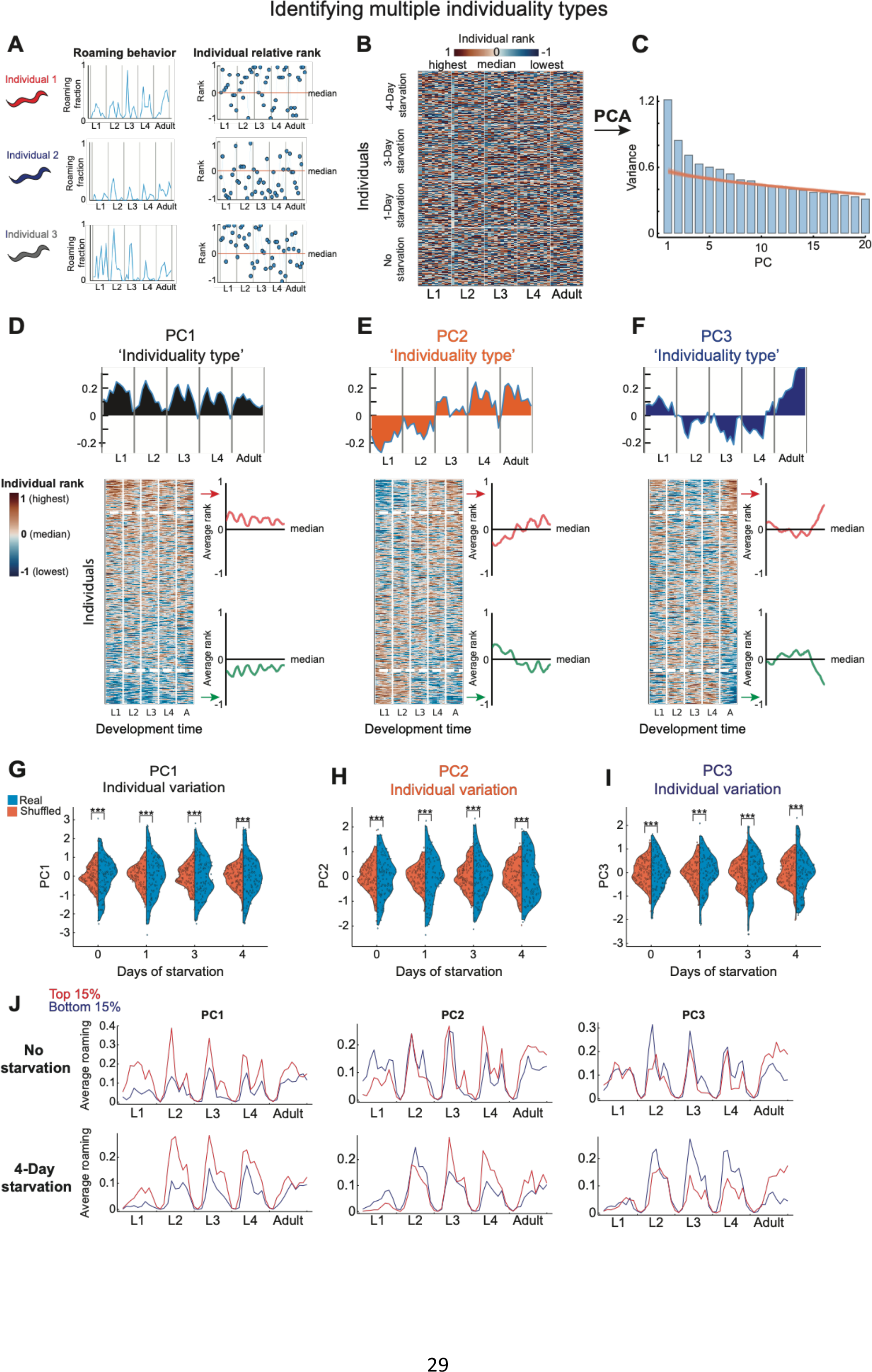
Unsupervised analysis of temporal individuality types across development. **(A)** Individual animals are ranked based on their roaming activity in each time-window compared to other individuals within the same experiment. **(B)** Heat-map represents relative rank of all N2 wild-type individuals (n=456) across 50 time windows (10 per developmental stage). **(C)** Variance explained by each of the first 20 PCs following PCA (blue bars), compared to the variance explained by the first 20 PCs of a shuffled dataset (500 repetitions, orange lines). **(D-F)** The first 3 PCs (PC1-PC3) represent the 3 temporal individuality types which explain the largest fraction of the variance in the individual rank dataset. For each PC individuality type shown are its components in each time window (top) and individuals sorted based on their PC score (bottom). Heat map is smoothed (4-bins window) for visual clarity. Average relative rank is plotted for extreme individuals (top and bottom 15%) within each individuality type. Midline represents the population median. **(G-I)** Distributions of individual scores (blue) within starved and unstarved wild-type populations for PC1-3 individuality types, compared to a shuffled dataset (orange). P-values were calculated using bootstrapping (see Methods). **(J)** Average roaming activity of top (red) and bottom (blue) 15% of extreme individuals within each of the PC1-3 individuality types in 4-day starved and unstarved wild-type populations. ***P<0.001.

We found that among the significant PC dimensions, the three major PCs (PC1-PC3) captured three distinct types of temporal individuality patterns within stressed and unstressed populations (Fig. 2D-F). PC1, which explained the majority of temporal variation in individual biases over time had eigenvector components of the same sign, indicating an individuality type of animals that consistently roam more or less than the population homogeneously across all developmental stages (Fig. 2D). The individuality type identified by PC1 unbiasedly recaptured a known mode of consistent individuality that was previously identified using a pre-defined index of long-term behavioral consistency across development (Stern et al., 2017) (Fig. S2B-C). This was further verified by the high correlation between the pre-defined consistency-index and scores of PC1 across individuals (R=0.9) (Fig. S2D). Interestingly, in contrast to PC1, PC2 and PC3 identified uncharacterized individuality types. PC2, which had opposite signs of eigenvector components before and after mid-development captured an individuality type that includes individuals that switch their behavioral bias once, during the L3 stage, from roaming more to roaming less than the population and vice versa (Fig. 2E). In addition, PC3, which had signs of eigenvector components that switch twice during development (at the end of L1 and L4), identified individuals that show the same behavioral bias during L1 and adulthood, which is opposite to their behavioral bias during intermediate stages (Fig. 2F). Both PC2 and PC3 scores of individuals did not correlate with the pre-defined consistency index (R=0.04 and 0.09, respectively) (Fig. S2D), further indicating that they indeed represent uncharacterized modes of temporal individuality types.

Inter-individual variation in a specific individuality type reflects how extreme individuals are within a population towards the identified individuality dimension. We found that wild-type populations with different early-life experiences show significantly extreme inter-individual variation in all PC1-3 individuality types, compared to a randomly shuffled rank dataset (p<0.001) (Fig. 2G-J; Fig. S2E), indicating the coexistence of these individuality types across stressed and unstressed individuals. Altogether, these results demonstrate the use of unsupervised analysis for identifying multiple individuality types across development and suggest a broad individuality space within isogenic populations.

### Dopamine buffers behavioral responses to early stress during intermediate developmental stages

Neuromodulatory pathways are known to establish internal behavioral states and modify them based on the environmental context (Harris-Warrick and Marder, 1991; Bargmann, 2012; Marder, 2012; Kennedy et al., 2014; Taghert and Nitabach, 2012; Nusbaum and Blitz, 2012). In particular, the bioamine dopamine was implicated in controlling a wide array of behavioral outputs at various timescales, ranging from minutes and hours, to long-term behavioral patterns that are regulated across life-stages (Marella et al., 2012; Omura et al., 2012; Sawin et al., 2000; Cermak et al., 2020; Stern et al., 2017). In *C. elegans*, dopamine is produced in a specific set of neuronal sites and its effects are known to be mediated by dopamine receptors that are localized to responding neurons (Sulston et al., 1975; Lints and Emmons, 1999; Chase et al., 2004; Tsalik et al., 2003).

To ask if dopamine acts across different developmental stages to shape the discontinuous pattern of long-term behavioral responses to early stress and to dissect its temporal requirement, we tracked the behavior of dopamine deficient *cat-2* animals following exposure to L1 starvation (Fig. 3A; Fig. S3A). When continuously grown in a food environment, *cat-2* individuals show a long-term roaming activity pattern that is similar to the wild-type population (Stern et al., 2017) (Fig. 1,3). However, we found that in contrast to stressed wild-type individuals that show buffering of behavioral responses during the L2, L3 and L4 intermediate stages, *cat-2* individuals that were exposed to early-starvation show reduction in overall roaming activity across all developmental stages, including during the intermediate stages (Fig. 3B,C). The behavioral effects of early starvation during mid-development in *cat-2* individuals were not only restricted to animals that were exposed to long starvation periods, as 1 day of early starvation was sufficient to induce a strong reduction in roaming activity during the L2-L4 intermediate stages (Fig. 3D; Fig. S3B). Interestingly, during the intermediate stages the effects on roaming activity were more pronounced during the first half of the stage, indicating within-stage regulation of behavioral response by dopamine (Fig. 3B,C). However, behavioral effects during L1 and adulthood following early stress were similar in *cat-2* and wild-type (Fig. 3B-D), implying that dopamine function is mainly required during intermediate developmental stages to buffer alterations in roaming activity in response to a transient early stress.

**Figure 3.**
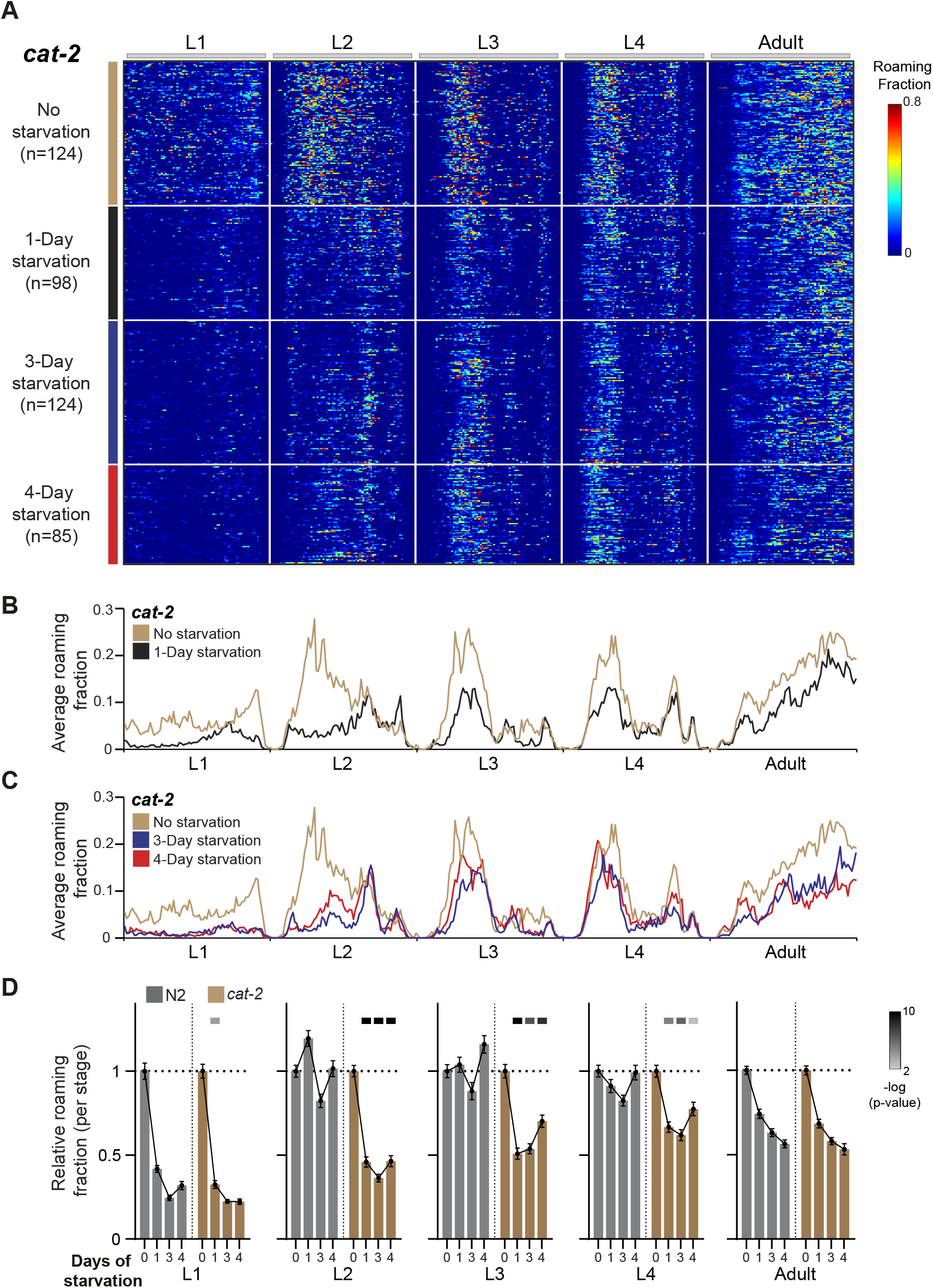
Dopamine buffers long-term behavioral effects during intermediate stages of development. **(A)** Roaming and dwelling behavior of *cat-2* animals without early starvation (n=124) and following 1-day (n=98), 3-day (n=124) and 4-day starvation (n=85). Each row indicates the age-normalized behavior of one individual across all developmental stages. The different stages are separated by white lines indicating the middle of the lethargus state. Color bar represents the fraction of time spent roaming in each of the 375 time-bins. **(B)** Average roaming fraction of 1-day starved *cat-2* animals compared to the unstarved population. **(C)** Average roaming fraction of 3- and 4-day starved *cat-2* animals compared to the unstarved population. **(D)** Average roaming fraction relative to the unstarved population in *cat-2* and wild-type individuals, in each developmental stage. Error bar indicates standard error of the mean. Upper bars indicate statistical significance (Wilcoxon rank-sum test, FDR corrected) of the difference in behavioral effect following early stress between the *cat-2* and wild-type populations (-log(P-value), indicated are p-values<0.01).

It was previously shown that during L2 to adulthood, *cat-2* individuals have higher instantaneous speed during roaming episodes (Stern et al., 2017; Sawin et al., 2000). We found that unlike stressed wild-type individuals in which roaming speed was decreased mainly during L1 and adulthood (Fig. S1E), *cat-2* mutants show lower speed also across the L2 and L3 stages following stress (Figure S3C).

To further ask if dopamine supplementation can restore the buffering of behavioral responses to stress during intermediate developmental stages, we exposed *cat-2* individuals to exogenous dopamine via their food after experiencing early starvation (Fig. 4A,B; Fig. S4A,B). We found that supplementing dopamine post-starvation was sufficient to recapitulate the buffering of only the roaming responses to stress in *cat-2* individuals during the L2,L3 and L4 stages (Fig. 4A,B; Fig. S4C,D). In contrast, exogenous dopamine did not restore roaming activity during L1 and adulthood (Fig. 4A,B; Fig. S4C). Overall, these results show that following early and transient starvation, dopamine acts to restrict long-term behavioral alterations in roaming activity, specifically during intermediate developmental windows.

**Figure 4.**
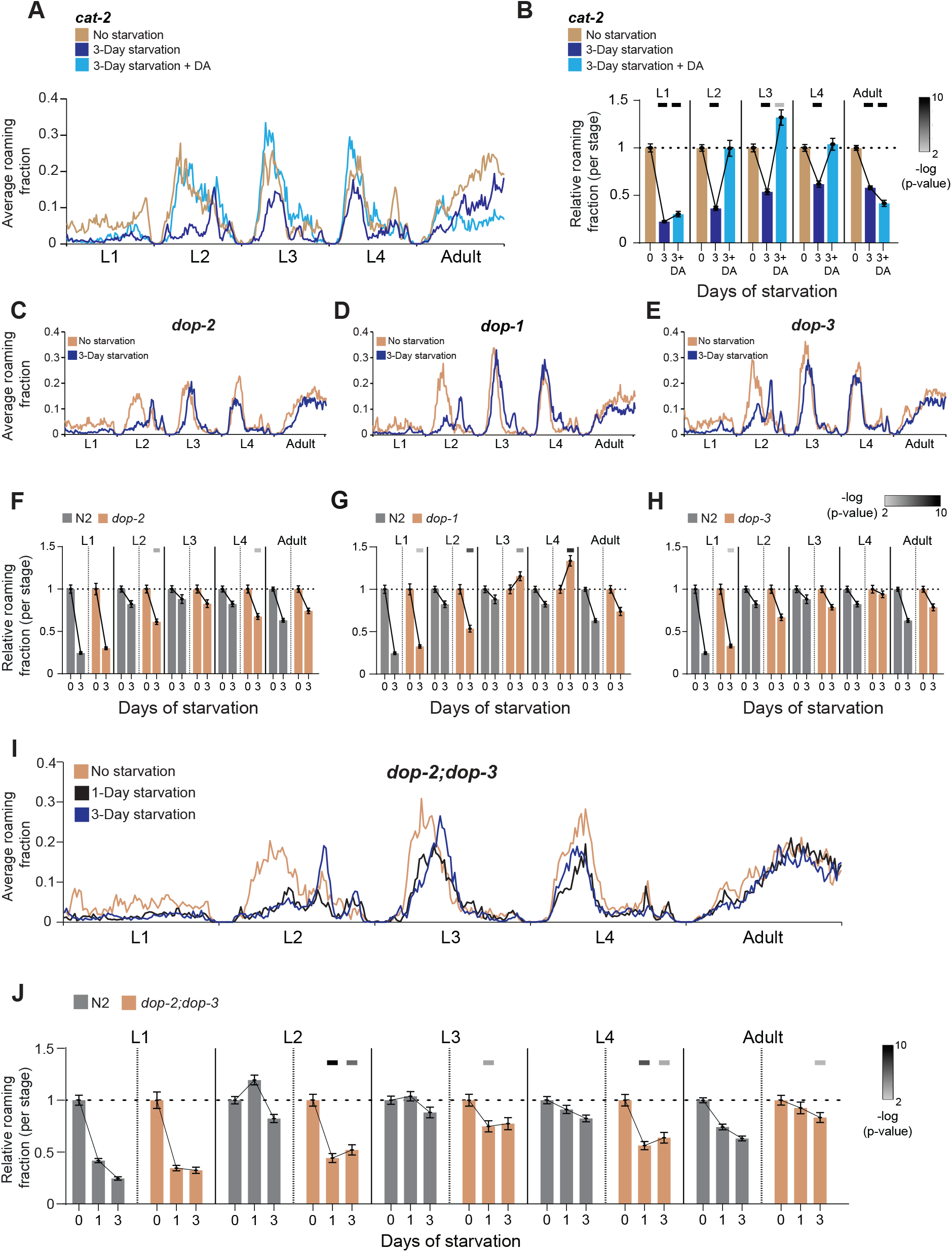
Effects of exogenous dopamine and temporally restricted functions of dopamine receptors across intermediate developmental stages. **(A)** Average roaming fraction of unstarved (n=124), 3-day starved (n=124) and 3-day starved with exogenous DA (n=63) *cat-2* populations. **(B)** Average roaming fraction relative to the unstarved population in 3-day starved and 3-day starved with exogenous DA *cat-2* individuals, in each developmental stage. Upper bars indicate statistical significance (Wilcoxon rank-sum test, FDR corrected) of difference in average roaming fraction compared to the unstarved population. **(C)** Average roaming fraction of 3-day starved (n=145) and unstarved (n=111) *dop-2* populations. **(D)** Average roaming fraction of 3-day starved (n=134) and unstarved (n=73) *dop-1* populations. **(E)** Average roaming fraction of 3-day starved (n=95) and unstarved (n=82) *dop-3* populations. **(F)** Average roaming fraction relative to the unstarved population in *dop-2* and wild-type individuals, in each developmental stage. **(G)** Average roaming fraction relative to the unstarved population in *dop-1* and wild-type individuals, in each developmental stage. **(H)** Average roaming fraction relative to the unstarved population in *dop-3* and wild-type individuals, in each developmental stage. **(I)** Average roaming fraction of 1-day starved (n=62), 3-day starved (n=63) and unstarved (n=69) *dop-2;dop-3* populations. **(J)** Average roaming fraction relative to the unstarved population in *dop-2;dop-3* and wild-type individuals, in each developmental stage. Upper bars in (F,G,H,J) indicate statistical significance (Wilcoxon rank-sum test, FDR corrected) of the difference in behavioral effect following early stress between the dopamine receptors mutants and N2 populations (-log(P-value), indicated are p-values<0.01).

### Specific dopamine receptors function during mid-development to mediate buffering of long-term behavioral responses

The buffering of behavioral effects during the L2, L3 and L4 intermediate developmental stages by dopamine led us to explore the temporal contribution of specific dopamine receptors during these development times. The *C. elegans* dopamine receptor DOP-1 is a D1-like receptor which signal through Gα_s/olf_ to activate adenylyl cyclase and DOP-2 and DOP-3 receptors are D2-like receptors which signal via Gα_i_ to suppress adenylyl cyclase (Chase et al., 2004; Sanyal et al., 2004; Sugiura et al., 2005; Suo et al., 2003).

To study the independent function of dopamine receptors we analyzed the long-term behavioral effects of early starvation in animals mutant for each of the single dopamine receptors. These analyses showed that each receptor has a different temporal effect on behavioral responses within the intermediate L2-L4 stages (Fig. 4C-H; Fig. S5). In particular, following 3 days of early starvation, *dop-2* animals showed strong roaming decrease during the L2 and L4 stages, but not during the L3 stage, compared to wild-type (Fig. 4C,F; Fig. S5). Interestingly, similar to *cat-2* mutants, roaming was mainly decreased in *dop-2* individuals during the first half of L2 and L4. In addition, *dop-1* individuals showed a roaming decrease during the L2 stage and opposite effects during L3 and L4 (Fig. 4D,G; Fig. S5) and *dop-3* animals showed weaker overall roaming response during the L2 stage (Fig. 4E,H; Fig. S5).

Previously, DOP-2 and DOP-3 were shown to cooperatively function to mediate behavioral effects of dopamine (Cermak et al., 2020; Suo et al., 2009). Therefore, we sought to test if simultaneous alteration of both dopamine receptors will recapitulate the full long-term behavioral effect during intermediate developmental stages, as shown in *cat-2* mutants. We found that following early starvation, *dop-2*;*dop-3* double mutants showed a decreased roaming activity across all intermediate stages (Fig. 4I,J; Fig. S6). These results imply that a deficiency in both DOP-2 and DOP-3 receptors is sufficient to recapitulate the behavioral effects in dopamine-deficient individuals during mid-development and suggest that buffering of long-term behavioral responses by dopamine is temporally regulated by the modular function of specific dopamine receptors.

### Serotonin promotes behavioral responses to early stress during early and late developmental stages

To ask if the stage-specific effects of early-life stress on developmental patterns of behavior are an integration of multiple temporal responses that are mediated by different neuromodulators, we also examined serotonin function in shaping long-term behavior following early starvation (Fig. 5A; Fig. S7A). Under normal growth conditions, serotonin-deficient *tph-1* individuals roam more than wild-type across all developmental stages (Flavell et al., 2013; Stern et al., 2017). We found that contrary to the effects of dopamine on the buffering of behavioral responses during intermediate stages, *tph-1* individuals that were exposed to 1 day of early starvation maintained their roaming activity during L1 and adulthood (Fig. 5B,D; Fig. S7B), compared to the strong roaming decrease generated in the wild-type population during these early and late stages. In addition, no significant difference in roaming response was shown during the L2-L4 intermediate developmental stages in the *tph-1* population following 1 day of early starvation (Fig. 5B,D; Fig. S7B), compared to wild-type.

**Figure 5.**
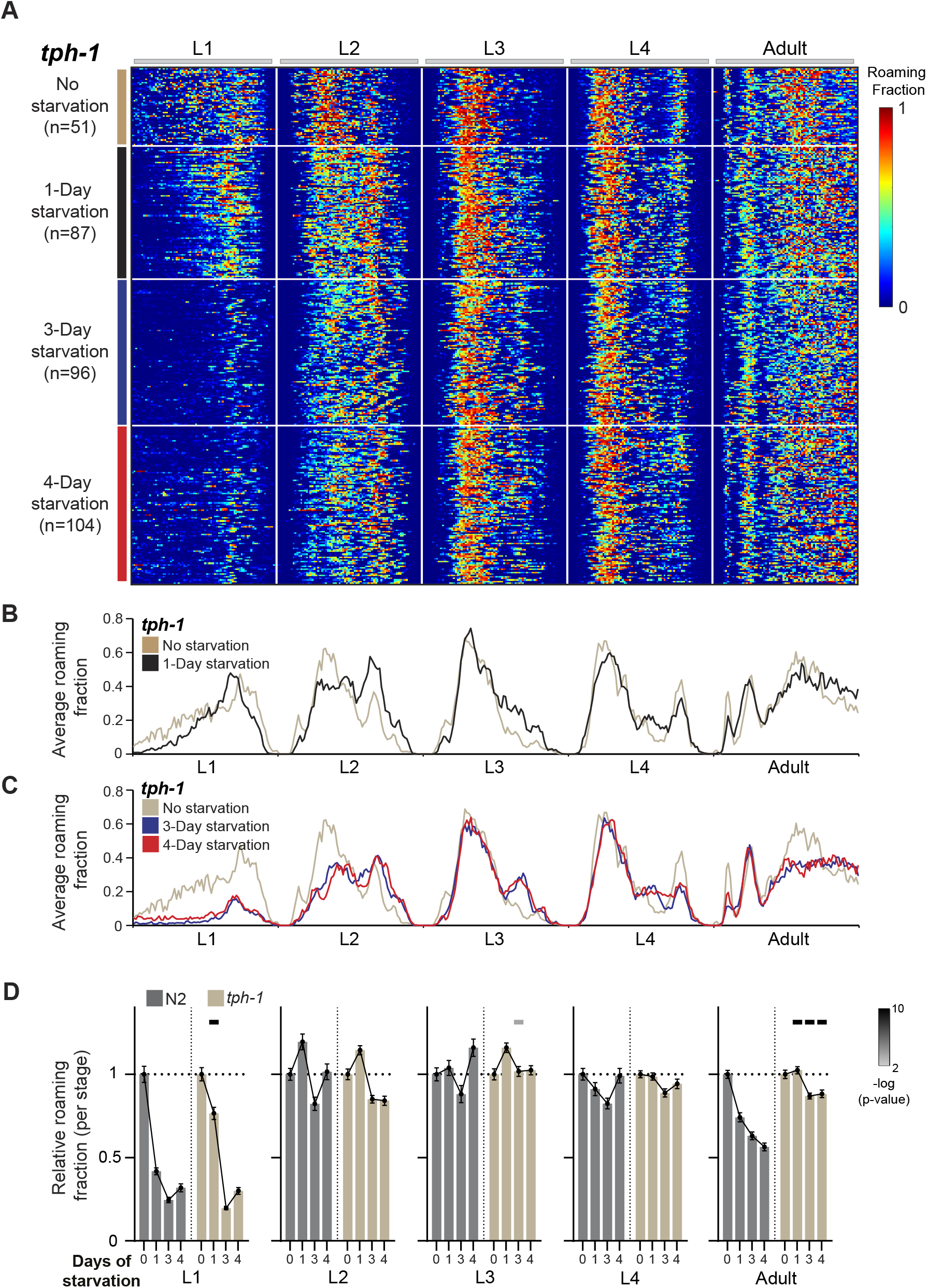
Serotonin is required for behavioral response during early and late developmental stages. **(A)** Roaming and dwelling behavior of *tph-1* animals without early starvation (n=51) and following 1-day (n=87), 3-day (n=96) and 4-day starvation (n=104). Each row indicates the age-normalized behavior of one individual across all developmental stages. The different stages are separated by white lines indicating the middle of the lethargus state. Color bar represents the fraction of time spent roaming in each of the 375 time-bins. **(B)** Average roaming fraction of 1-day starved *tph-1* animals compared to the unstarved population. **(C)** Average roaming fraction of 3- and 4-day starved *tph-1* animals compared to the unstarved population. **(D)** Average roaming fraction relative to the unstarved population in *tph-1* and wild-type individuals, in each developmental stage. Error bar indicates standard error of the mean. Upper bars indicate statistical significance (Wilcoxon rank-sum test, FDR corrected) of the difference in behavioral effect following early stress between the *tph-1* and N2 populations (-log(P-value), indicated are p-values<0.01).

To test if longer starvation periods early in life will establish behavioral effects during L1 and adulthood we further exposed *tph-1* animals to 3- and 4-days of early-starvation. We found that long starvation periods led to a reduction in roaming activity in the L1 stage of *tph-1* animals. However, during the adult stage, *tph-1* individuals were still less responsive to early stress, compared to the strong decrease in roaming in the wild-type (Fig. 5C,D; Fig. S7B). In addition, behavioral responses to early stress were similar in *tph-1* and wild-type individuals during the intermediate L2-L4 stages, indicating that serotonin effects are specific to shaping behavioral responses during L1 and adulthood.

These results show that dopamine and serotonin functions are opposite and segregated across developmental stages in regulating long-term roaming behavior following stress. While dopamine buffers behavioral modifications during intermediate stages of development, serotonin functions to promote behavioral sensitivity to early starvation during the early L1 stage and adulthood. Interestingly, functional segregation among dopamine and serotonin regulation is behavior-specific, as the long-term effects of early stress on roaming speed were similar in *tph-1* and *cat-2* individuals (Fig. S7C;Fig. S3C).

### Early-life experiences and neuromodulation shape variation in individuality types within populations

To ask if neuromodulatory pathways affect specific individuality types to reshape inter-individual variation within stressed and unstressed populations, we performed the PCA on pooled data across the wild-type, *cat-2* and *tph-1* populations (Fig. S8A). This analysis identified highly similar PC1-3 individuality types to the ones obtained by analyzing only wild-type individuals (Fig. 6A-C; fig. 2D-F). It was previously shown that *tph-1* individuals lacking serotonin show reduced levels of consistent individual biases across development, measured by the pre-defined consistency index (Stern et al., 2017). Similarly, we found that inter-individual variation in PC1 individuality type which reflects an individuality mode of consistent homogeneous bias across all developmental stages (Fig. S8B-D) was reduced within the unstressed and 1-day starved *tph-1* populations, compared to wild-type (Fig. 6D). However, we found that following long starvation periods (3 and 4 days), inter-individual variation in PC1 type was not significantly different in *tph-1* individuals, compared to the wild-type population (Fig. 6D). The increase in PC1 inter-individual variation in *tph-1* individuals following stress indicates that early starvation experiences may generate extreme behavioral consistency in a specific neuromodulatory context where consistency levels are initially low. In contrast to *tph-1* mutants, dopamine-deficient *cat-2* individuals did not show a significant difference in PC1 inter-individual variation across all conditions, compared to wild-type (Fig. 6D).

**Figure 6.**
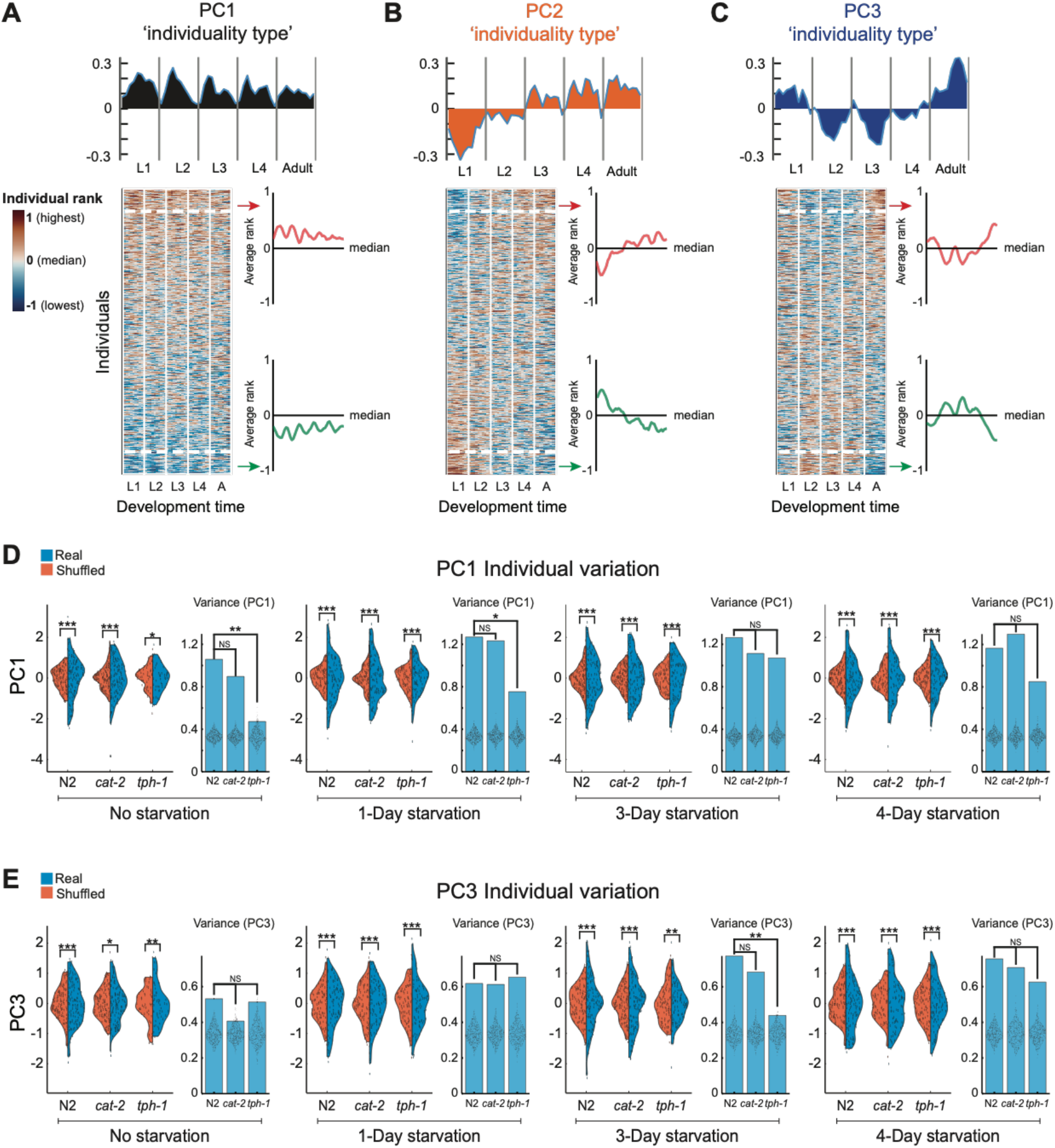
Neuromodulatory and experience-dependent effects on individuality types within stressed and unstressed populations. **(A-C)** The first 3 PCs (PC1-PC3) represent the 3 temporal individuality types which explain the largest fraction of the variance in the pooled individual rank dataset of the wild-type, *cat-2* and *tph-1* populations. For each PC individuality type shown are its components in each time window (top) and individuals sorted based on their PC score (bottom). Heat map is smoothed (4-bins window) for visual clarity. Average relative rank is plotted for extreme individuals (top and bottom 15%) within each individuality type. Midline represents the population median. **(D)** Distributions of PC1 individual scores (blue) within starved and unstarved wild-type and mutant populations, compared to shuffled dataset (orange). Bar plots represent inter-individual variation in PC1 individual scores. Each dot within bars represents PC1 variation in individual scores within a shuffled dataset (500 repetitions). **(E)** Distributions of PC3 individual scores (blue) within starved and unstarved wild-type and mutant populations, compared to shuffled dataset (orange). Bar plots represent inter-individual variation in PC3 individual scores. Each dot within bars represents PC3 variation in individual scores within a shuffled dataset (500 repetitions). P-values in D,E were calculated using bootstrapping (see Methods). *P<0.05, ** P<0.01, ***P<0.001.

In addition, we found that serotonin also alters the effects of stress on inter-individual variation in PC3 type (double bias switching across development). In particular, while the wild-type population showed an increased inter-individual variation in PC3 type following 3 days of early starvation, *tph-1* mutants had significantly lower PC3 inter-individual variation (Fig. 6E) following the same stressful experience. The effect of serotonin on PC3 inter-individual variation was specific, as dopamine-deficient *cat-2* individuals showed similar increase in PC3 inter-individual variation following 3 days of starvation when compared to wild type (Fig. 6E). Interestingly, neuromodulation affected only a fraction of the identified individuality types. Inter-individual variation in PC2 type (single bias switching during mid-development) was not significantly different in *tph-1* and *cat-2* mutants when compared to wild type populations (Fig S8E). Overall, these results imply that inter-individual variation in a spectrum of individuality types may be dynamically structured by the integration of the population’s neuromodulatory state and its early life-experience.

## Discussion

Spontaneous behavioral patterns across development are structured in time and shaped by the integration of the individual’s internal state and its past and current environments. In this work we studied how developmental patterns of behavior and inter-individual variation are dynamically affected by early-life starvation and the neuromodulatory pathways that organize these long-term behavioral responses. The effects of transient early experiences on neuronal and behavioral states during specific developmental stages were studied across species (Horn, 1998; Kimmel et al., 1974; Nakamori et al., 2013; Pradhan et al., 2019; Remy and Hobert, 2005; Wilson and Sullivan, 1994). However, how transient environmental experiences early in development continuously reshape behavior throughout the full developmental trajectory of the organism is unknown.

Here, we utilized long-term behavioral tracking systems at high spatiotemporal resolution (Stern et al., 2017) to analyze and compare long-term alterations of behavioral patterns across and within all developmental stages of *C. elegans*, following transient periods of starvation early in life. Our results show that early starvation generates stage-specific behavioral responses that are discontinuous across development, manifested by stronger decrease in roaming activity during early and late stages, compared to intermediate developmental stages. These variable influences of early starvation across development time suggest that while the memory of early experiences is maintained to adulthood, behavioral changes are buffered during mid-development.

As imprinting of early memories was shown to have an adaptive value for later stages of life (Immelmann, 1975), we hypothesized that neuronal mechanisms actively buffer behavioral alterations at specific development times so as to support the exploratory activity of individuals during critical developmental windows. Building on this idea, we further analyzed the contribution of neuromodulatory pathways for shaping the stage-specific patterns of behavioral responses across development. Neurotransmitters and hormones were shown to regulate behavioral patterns across development (Sisk and Foster, 2004; Truman, 2005; Wigglesworth, 1936; Aton et al., 2005; Park and Hall, 1998; Rehm et al., 2008). In *C. elegans* populations grown continuously on food, neuromodulators show both consistent and time-dependent behavioral effects at specific developmental windows (Stern et al., 2017). We found that following early transient stress, dopamine and serotonin control of long-term behavioral responses is opposite and temporally segregated over development time. Dopamine was required for behavioral buffering during intermediate developmental stages and serotonin established behavioral responses to early stress during early and late developmental stages. In *C. elegans,* dopamine is produced in four pairs of neurons: CEPV, CEPD, ADE, and PDE (Sulston et al., 1975; Lints and Emmons, 1999) and was shown to be required for controlling locomotory patterns (Omura et al., 2012) and coupling of behavioral programs (Cermak et al., 2020). In particular, dopamine was shown to decrease the instantaneous speed of worms grown on food, compared to non-food environment (Sawin et al., 2000). We found that following early-life starvation, dopamine is required for buffering roaming decrease during intermediate developmental stages. These diverse behavioral effects on different locomotory parameters suggest that dopamine function is variable under different environmental contexts and at different timescales.

By analyzing single dopamine receptors (DOP-1, DOP-2 and DOP-3), we found that their functions are differentially distributed during intermediate developmental stages. We further found that both DOP-2 and DOP-3 function (D2-like receptors) are required for establishing behavioral buffering across all intermediate developmental stages (L2-L4). The modular regulation by each of the receptors and their cooperative function imply that dopamine receptors’ temporal requirement may be super-imposed in time to mediate buffering of behavioral responses at specific developmental windows. Interestingly, the expression patterns of the three dopamine receptors within the *C. elegans* nervous system are partially overlapping (Tsalik et al., 2003; Chase et al., 2004; Sanyal et al., 2004; Suo et al., 2003), raising the possibility that different subnetworks within the nervous system function to temporally regulate behavioral buffering across development. Similarly, the function of serotonin receptors in maintaining patterns of roaming activity in unstressed individuals was also shown to be modular across developmental stages (Stern et al., 2017), suggesting a common principle of temporal regulation of behavior by neuromodulatory receptors.

In contrast to dopamine function during intermediate developmental stages, we showed that serotonin promotes roaming responses to early stress during L1 and adulthood. Under normal growth conditions, serotonin is known to regulate roaming behavior in *C. elegans* across all developmental stages, (Flavell et al., 2013; Stern et al., 2017) and is required for long- and short-term associative olfactory memory (Jin et al., 2016; Zhang et al., 2005). In rodents, serotonin and dopamine interact to establish motor patterns (Sasaki-Adams, 2001). It is plausible that the complexity of long-term behavioral responses to early-stress reflects a time-integration of the function of multiple neuromodulators, each of them acting at different development times and with different intensity.

Long-term individuality in behavior is observed across species, even within genetically-identical populations that were raised in the same environment (Buchanan et al., 2015; Freund et al., 2013; Stern et al., 2017; Honegger et al., 2020; Kain et al., 2012; Linneweber et al., 2020; Schuett et al., 2011, 2011). However, the effects of early stressful experiences on patterns of individual biases within isogenic populations are less explored. The long-term behavioral tracking of single animals allowed us to ask how early-life stress modifies patterns of inter-individual behavioral variation and whether neuromodulation control the structure of individuality under stress. Individuality is classically defined as the tendency of an individual to show the same behavioral bias relative to the population over long timescales. We hypothesized that within isogenic populations, individuals may show alternative modes of temporal behavioral biases across development that are not random and represent alternative individuality types.

By using an unbiased approach of dimensionality reduction, we found multiple individuality types that coexist within stressed and unstressed populations. While the main PC1 individuality type recaptured a known individuality type of consistent individual biases over time (Stern et al., 2017), PC2 and PC3 identified alternative individuality types that are significant within populations and represent individuals that show switching of behavioral bias, relative to the population, at specific developmental times. These results further extend the view of long-term behavioral individuality, implying a wide spectrum of alternative individual biases within populations (Tang et al., 2012). A plausible explanation for the coexistence of multiple individuality types is that, upon stress or another unpredictable environment, it will be beneficial for the population to dynamically reshape the composition of individuality types so as to modify individual strategies and increase the population’s chance of survival (Cooper and Kaplan, 1982; Honegger and de Bivort, 2018).

Neuromodulation was previously shown to affect levels of consistent individual biases (Honegger et al., 2020; Kain et al., 2012; Pantoja et al., 2016; Stern et al., 2017). We tested how early-life experiences and neuromodulation shape the identified individuality types across development. Interestingly, we found that inter-individual variation in specific types depends on both the neuromodulatory state of the population and its early experience. An open question is what are the sources of variation within the nervous system that give rise to different individuality types? Underlying differences among individuals may include diversity in gene-expression patterns (Casanueva et al., 2012), nervous system structure (Witvliet et al., 2021; Brittin et al., 2021; Linneweber et al., 2020; Churgin et al., 2021), and underlying persistent differences in neruomodulatory parameters that have phenotypic effects under extreme conditions (Marder et al., 2022). It is plausible that some of this variation, which is partially stochastic by nature, may generate different behavioral biases of individuals within isogenic populations.

The behavioral patterns explored in this study represent only a small fraction of the behavioral space available to the organism (Ahamed et al., 2021; Anderson and Perona, 2014; Brown and de Bivort, 2018; Schwarz et al., 2015). We anticipate that an extended supervised and unsupervised behavioral classification across development will shed light on the overall reorganization of individuality types and the contribution of both internal neuronal states and external environments to the diversity in long-term behavioral structures within populations.

## Author contributions

R.A. and S.S. designed experiments. R.A conducted experiments. R.A. S.S. and Y.H. analyzed and interpreted data, and S.S, R.A and Y.H. wrote the paper.

## Acknowledgments

We thank the Caenorhabditis Genetics Center (CGC) for strains; Sagy Levi, Cori Bargmann and the members of our laboratory for comments on the manuscript. Some strains were provided by the CGC, which is funded by the NIH Office of Research Infrastructure Programs (P40 OD010440).This work was supported by the European Research Council ERC-2019-STG and the Israel Science Foundation grant 3035/20.

## Materials and Methods

### Growth conditions and starvation protocol

*C. elegans* worms were maintained on NGM agar plates, supplemented with E. coli OP50 bacteria as a food source. For behavioral tracking, we imaged single individuals grown in custom-made laser-cut multi-well plates. Each well (10mm diameter) was seeded with a specified amount of OP50 bacteria (10 uL of 1.5 OD) that was UV-killed before the experiment to prevent bacterial growth. For the starvation experiments, eggs were collected from isogenic populations using a standard bleaching protocol, into an agar plate without OP50 bacteria. Newly hatched L1 larvae were starved for a specified time-window (L1 arrest of 1, 3 or 4 days) before being transferred to the imaging multi-well plates. For tracking behavior without early L1 starvation, animals were monitored immediately after hatching in the multi-well plates. For post-starvation dopamine supplementation, 1000ug dopamine (Sigma) was added to 1 ml of OP50 solution.

### *C. elegans* strains

Strains used in this study:

Wild-type Bristol N2

MT15434 *tph-1* (mg280) II

CB1112 *cat-2* (e1112) II

LX645 *dop-1* (vs100) X

LX702 *dop-2* (vs105) V

LX703 *dop-3* (vs106) X

LX704 *dop-2* (vs105) V; *dop-3*(vs106) X

### Imaging system

Longitudinal behavioral imaging was performed using custom-made imaging systems. Each imaging system consists of an array of six 12 MP USB3 cameras (Pointgrey, Flea3) and 35 mm high-resolution objectives (Edmund optics) mounted on optical construction rails (Thorlabs). Each camera images up to six wells, each containing an individual grown in isolation. Movies are captured at 3 fps with a spatial resolution of ~9.5 um. For uniform illumination of the imaging plates we used identical LED backlights (Metaphase Technologies) and polarization sheets. To tightly control the environmental parameters during the experiment, imaging was conducted inside a custom-made environmental chamber in which temperature was controlled using a Peltier element (TE technologies, temperature fluctuations in the range of 22.5 ± 0.1°C). Humidity was held in the range of 50% +/− 5% with a sterile water reservoir and outside illumination was blocked, keeping the internal LED backlights as the only illumination source. Movies from the cameras were captured using commercial software (FlyCapture, Pointgrey) and saved on two computers (3 cameras per computer; each computer has at least 8-core Intel i7/i9 processor and 64 GB RAM).

### Imaging data processing for extracting locomotion trajectory

To extract behavioral trajectories of animals across the experiment, captured movies were analyzed by custom-made script programmed in MATLAB (Mathworks, version 2019b) (Stern et al., 2017). In each frame of the movie and for each behavioral arena, the worm is automatically detected as a moving object by background subtraction, and its XY position is logged (center of mass). In each experiment, 600,000-1,000,000 frames per individual are analyzed using ~50 processor cores in parallel to reconstruct the full behavioral trajectory of individuals over days of measurements across development. The total time of image processing was 3-7 days per experiment. Egg hatching time of each individual in the experiment is automatically marked by the time when activity can be detected in the behavioral arena. The middle of the lethargus periods, in which animals stop their locomotion and molt, were defined as the transition points between different stages of development (based on 10s time-scale speed trajectories over time, smoothed over 300 frames). To synchronize temporal behavioral trajectories of different individuals we age-normalized individuals by dividing the behavioral trajectory of each life stage into a fixed number of time windows.

### Behavioral parameters quantification

For each individual, we differentiate between roaming and dwelling states by averaging speed (μm/s) and angular velocity (absolute deg/s) over 10 seconds using a rolling time window, and generating a 2D probability distribution of these two behavioral parameters for all intervals in each time-bin along the experiment (50 × 50 bins distribution, speed bin size: 7.59 um/s, angular velocity bin size: 3.6 deg/s) (Stern et al., 2017). Drawing a diagonal through the probability distribution separated roaming and dwelling states, such that intervals in the distribution bins below the diagonal were classified as roaming intervals and intervals in bins above the diagonal were classified as dwelling intervals (Arous et al., 2009; Flavell et al., 2013; Stern et al., 2017). The behavior of each animal over time could be quantified as a sequence of roaming and dwelling intervals. The fraction of time spent roaming of the individual in a time bin represents the fraction of these intervals classified as roaming states within a given time bin. For each developmental stage, we examined the two-dimensional probability distribution of the whole population and changed the slope of the diagonal to classify roaming and dwelling appropriately (slopes: 5,2.5,2.3,2,1.5 for the L1,L2,L3,L4 and adult stages, respectively). Based on these roaming and dwelling classifications we further quantified the average instantaneous speed of the animal during roaming episodes (μm/s).

### Unsupervised quantification of temporal individuality types

#### Ranking and behavioral bias

Individuals within the population were ranked based on their roaming behavior in 50 time bins (10 per stage), relative to the population measured within the same experiment. More explicitly, within each experiment, individuals were ranked in each time bin by the fraction of time within the bin spent roaming. Ties were resolved as fractional ranks (“1 2.5 2.5 4 ranking”). This produces a rank *r*_*i,k*_ for the *i*th individual in the *k*th time bin, between 1 and *n*_*i*_, where *n*_*i*_ is the number of individuals measured in the experiment which includes individual *i*. These ranks were normalized to obtain *bias* values between −1 and 1, as 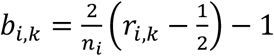. Thus, a bias *b*_*i,k*_ = 0 is obtained when the roaming fraction of worm *i* in bin *k* is the median roaming fraction for that experiment. A positive bias occurs in a time bin where a worm roams more than the median roaming fraction for that time bin across its experiment, and a negative bias where it roams less than the median. Particularly, in each time bin, the worms with the highest and lowest roaming fraction within an experiment have ranks (1 − 1/*n*) and (−1 + 1/*n*) respectively, where *n* is the number of worms in the experiment.

#### Identification of temporal bias patterns

To identify temporal individual biases that are dominant within the isogenic populations, we performed principal component analysis (PCA) on individuals’ biases across time bins.

This analysis represents each individual’s sequence of biases 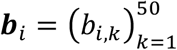, as a weighted sum of Principal Components (PCs),

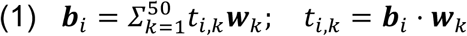

where ***w***_*k*_ is the *k*th PC, *t*_*i,k*_ the *k*th PC score for the *i* th individual. Note that equation (1) does not include a mean term, since the mean of all biases at each time bin is zero by construction. The first PC ***w***_1_ is the direction where the variance of the population is highest (namely, the unit vector for which the variance of the dot product ***b***_*i*_ ⋅ ***w***_1_ across the population is maximized). The second PC ***w***_2_ is the direction of highest variance in the subspace orthogonal to ***w***_1_, and so on. The PCs are obtained as eigenvectors of the covariance matrix of the input vectors ***b***_*i*_. When computing PCA across several experimental conditions, each individual bias vectors ***b***_*i*_ was weighted in inverse proportion to the number of individuals in the same condition (strain and starvation level), so that each condition has equal weight. Specifically, the variance which is maximized by the PCs is the weighted variance of the bias vectors, where ***b***_*i*_ is assigned the weight 1/*n*_*i*_. In practice, this is achieved by computing the PCs as eigenvectors of the *weighted* covariance matrix. Early principal components thus represent the temporal patterns of individual biases ***b***_*i*_ which account for the most variance in the rank sequences. Statistical significance of PCs was assessed by comparing the variances of PC scores generated from the real individual rank dataset to variances calculated from a randomly shuffled rank dataset (500 repetitions). This test identifies which PCs account for a higher fraction of variance than would be expected by chance in a population whose behavior has no underlying temporal correlations. Inter-individual variance in each PC score was calculated as a dispersal parameter of the population for each PC individuality type. To quantify significant differences in PC score inter-individual variance between conditions, we used a permutation test where individuals in each pair of conditions were randomly reassigned to two populations of the same size. Significance values were computed from 1000 such reassignments for each pair of conditions. The test was repeated multiple times to verify the robustness of the analysis.

#### Quantification of individual consistency index

Individuals within the population were ranked based on their behavior in 50 time bins (10 per stage). We then quantified the homogeneous consistent bias in the individual’s behavior relative to the population (Stern et al., 2017) by calculating for each individual the log_2_(number of time bins in which the individual’s roaming fraction is higher than the population median / number of time bins in which the individual’s roaming fraction is lower than the population median) (‘consistency index’). This measure gives positive values to individuals that tend to have positive bias (higher than median roaming fraction) across time, negative values to individuals that tend to have negative bias across time, and values close to 0 for individuals that do not show any bias toward higher or lower roaming fraction. Inter-individual variance in consistency index was calculated as a dispersal parameter of the population which indicates the overall consistent behavioral bias in the population. To quantify significant differences in inter-individual variance of behavioral consistency between conditions, we used a permutation test, as used for comparing variance of PC scores (see ‘Identification of temporal bias patterns’).

---

**Figure S1.**
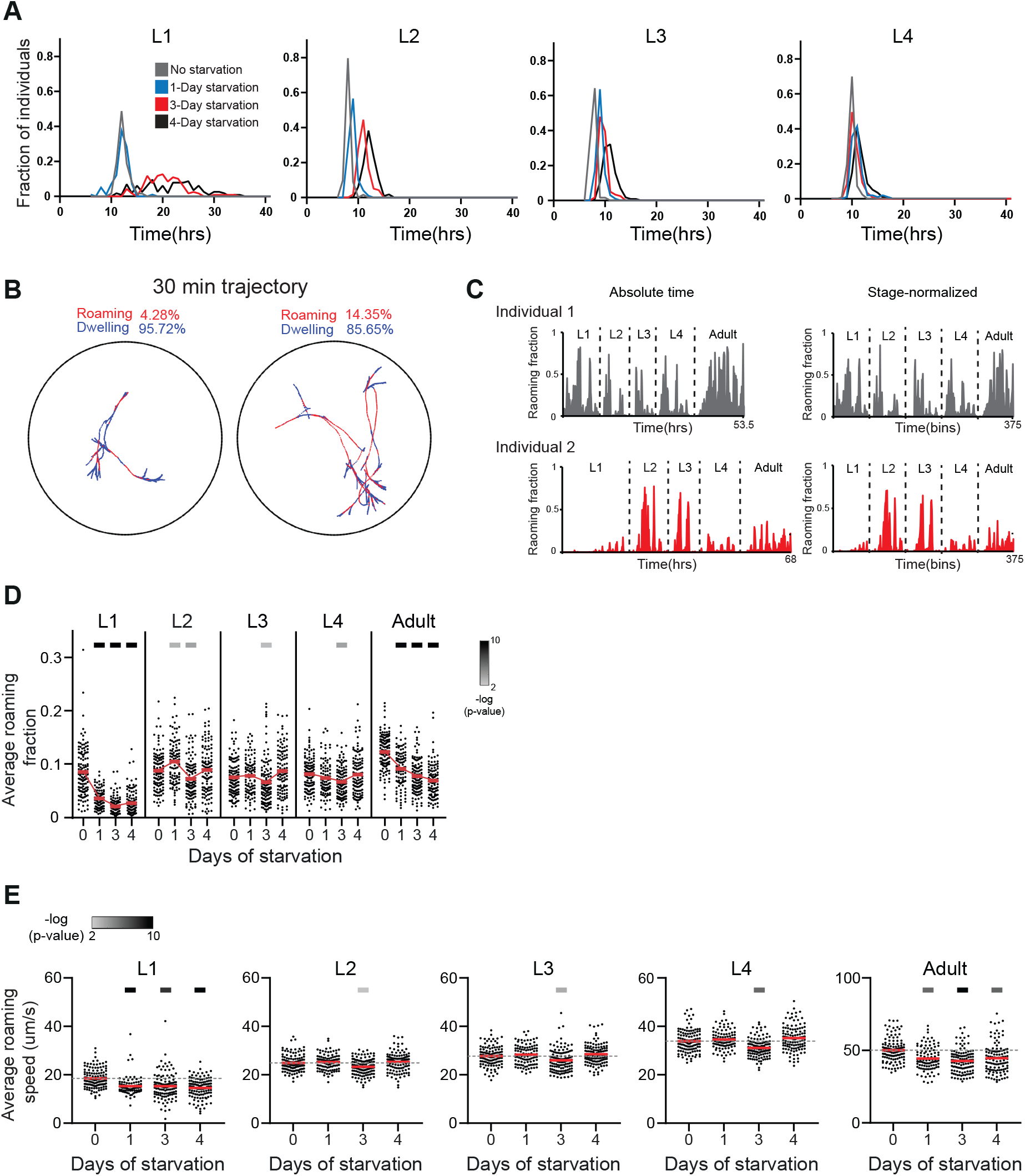
Development-time, behavioral trajectory synchronization and roaming quantification in starved and unstarved wild-type individuals. Related to Figure 1. **(A)** Distributions of development time across L1-L4 larval stages within starved and unstarved wild-type populations (No starvation n=123, 1-day starvation n=99, 3-day starvation n=119 ,4-day starvation n=115). **(B)** Examples of 30 minutes behavioral trajectories of adult individuals. Roaming and dwelling episodes are indicated in red and blue, respectively. **(C)** Absolute time (left) and age-normalized (right) roaming activity of a starved (red) and unstarved (gray) single individuals across all developmental windows. Normalization synchronizes behavioral trajectories of different individuals by dividing each life-stage of each individual into 75 equal time windows. **(D)** Average roaming fraction of starved and unstarved individuals in each developmental stage. Each point represents a single individual, red bars represent population mean. **(E)** Average roaming speed of starved and unstarved individuals in each developmental stage. Each point represents a single individual, red bars represent population mean. Upper bars in (D,E) indicate statistical significance (Wilcoxon rank-sum test ,FDR corrected) of the difference between unstarved and starved populations in each developmental stage (-log(P-value), indicated are P-values<0.01).

**Figure S2.**
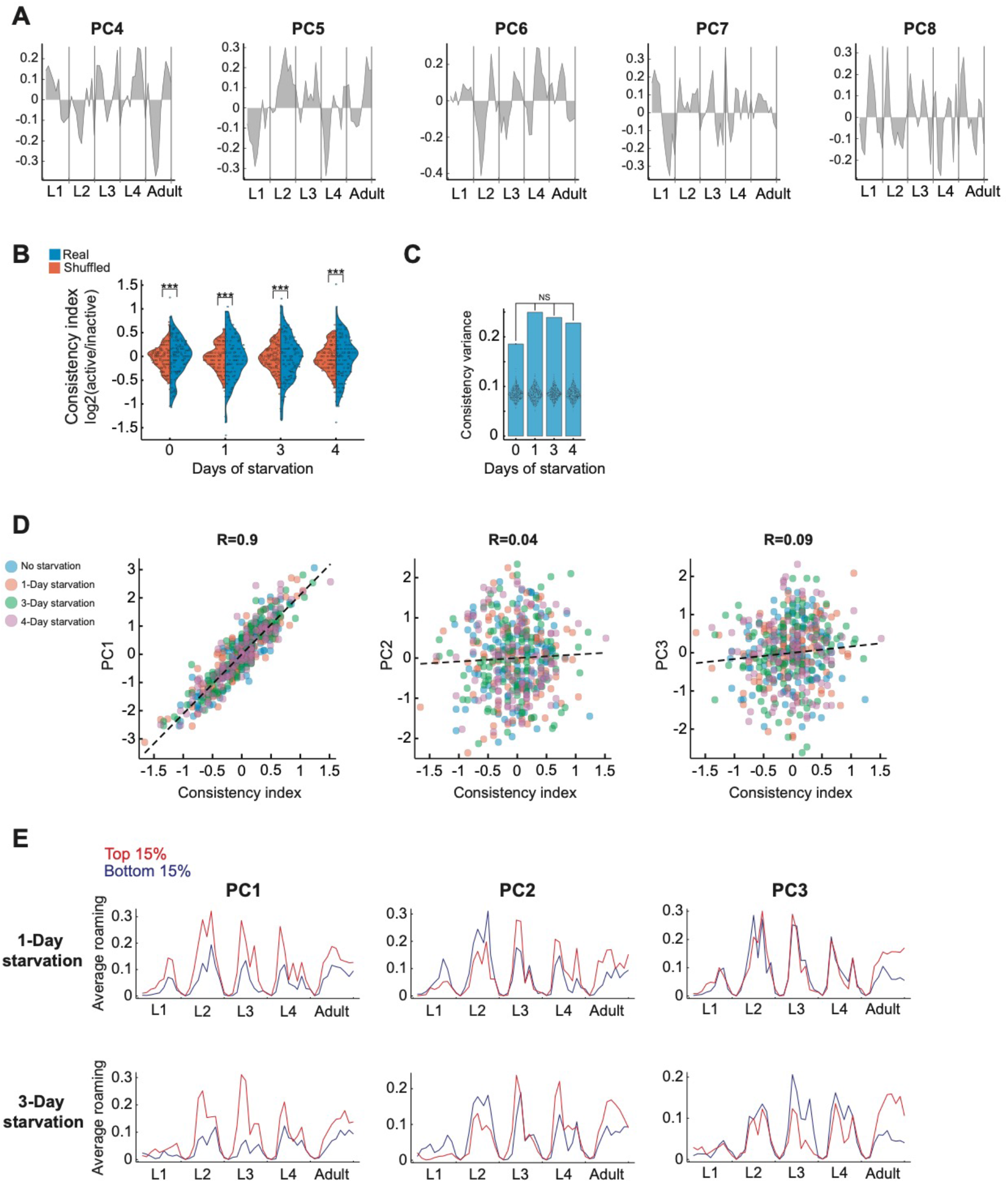
PCA and behavioral consistency analyses. Related to Figure 2. **(A)** PC4-8 vectors across developmental time-bins following PCA of wild-type behavioral rank dataset. The first 8 PCs are significant relative to a shuffled rank dataset (P<0.001). **(B)** Distributions of consistency indices representing homogeneous behavioral consistency of individuals across development in starved and unstarved wild-type individuals (blue) compared to shuffled dataset (orange). P-value was calculated using bootstrapping (see Methods). **(C)** Inter-individual variation in behavioral consistency indices in (B). P-values were calculated between variation values of starved and unstarved populations using bootstrapping (see Methods). Each dot within bars represents inter-individual variation within a shuffled dataset (500 repetitions). **(D)** Correlation between behavioral consistency indices and PC1-3 scores of individuals within stressed and unstressed wild-type populations. Each dot is a single individual, colored by starvation condition. Dotted line is linear least-squares regression with intercept. Pearson correlation coefficient R is noted above each subplot. **(E)** Average roaming activity of top (red) and bottom (blue) 15% of extreme individuals within each of the PC1-3 individuality types in 1-day and 3-day starved wild-type populations. ***P<0.001.

**Figure S3.**
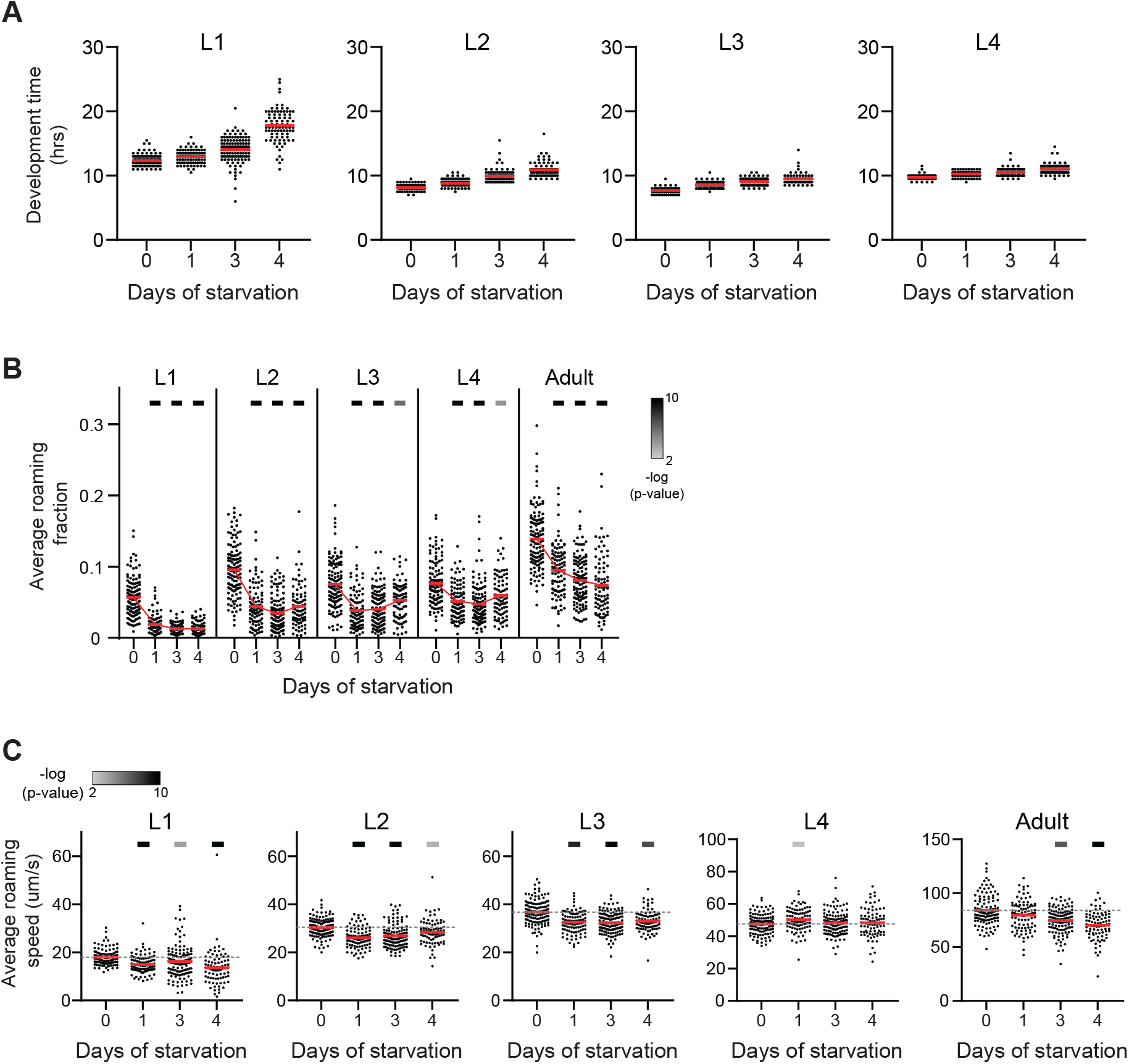
Development-time and roaming quantification in starved and unstarved *cat-2* individuals. Related to Figure 3. **(A)** Development time across L1-L4 larval stages within starved and unstarved *cat-2* populations (no starvation n=124; 1-day starvation n=98; 3-day starvation n=124; 4-day starvation n=85). **(B)** Average roaming fraction of starved and unstarved *cat-2* individuals in each developmental stage. Each point represents a single individual, red bars represent population mean. **(C)** Average roaming speed of starved and unstarved *cat-2* individuals in each developmental stage. Each point represents a single individual, red bars represent population mean. Upper bars in (B,C) indicate statistical significance (Wilcoxon rank-sum test ,FDR corrected) of the difference between unstarved and starved populations in each developmental stage (-log(P-value), indicated are P-values<0.01).

**Figure S4.**
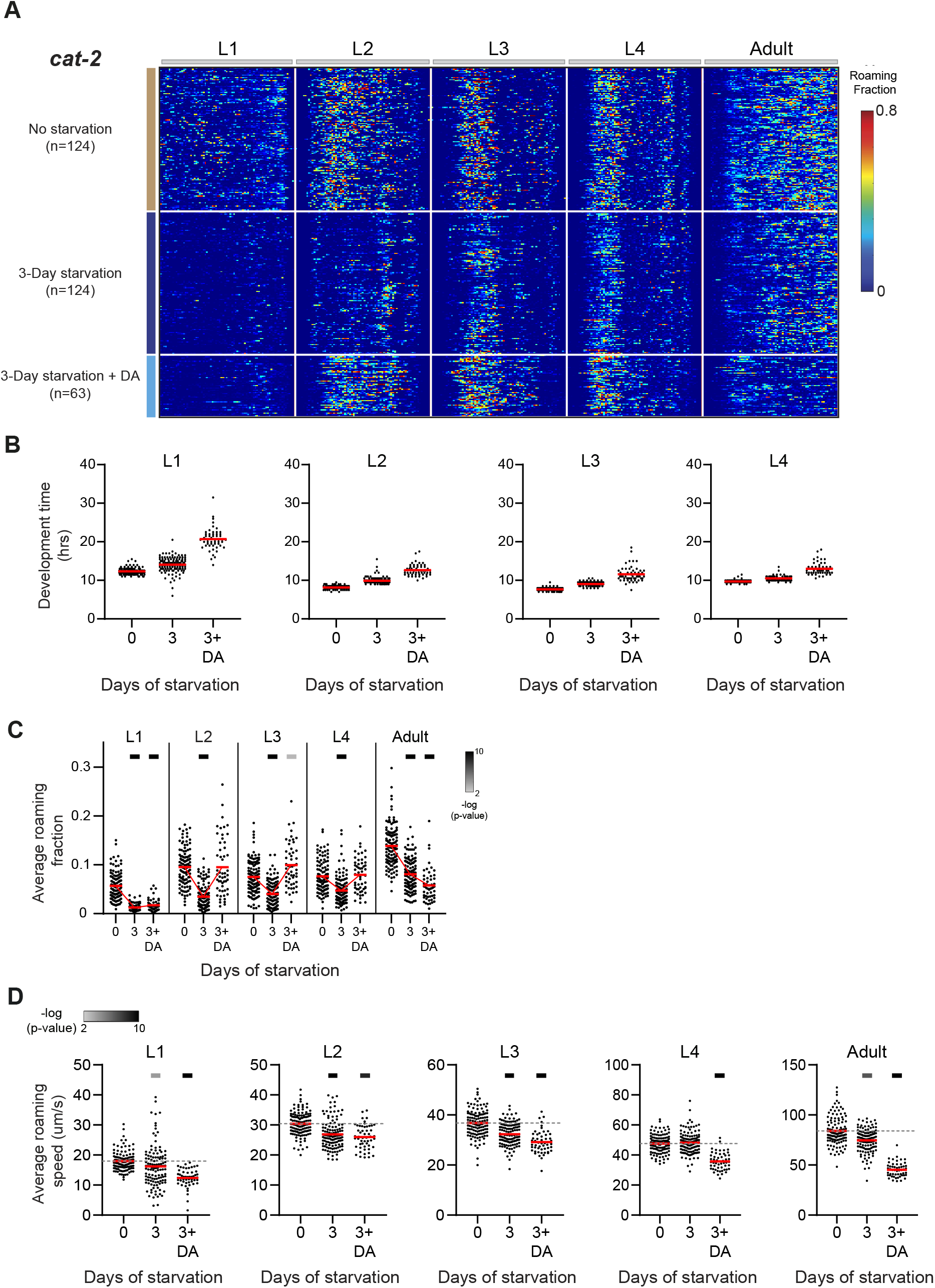
Development-time and roaming quantification in unstarved, starved and starved with exogenous DA *cat-2* mutants. Related to Figure 4. **(A)** Roaming and dwelling behavior of *cat-2* animals without early starvation (n=124), following 3 days of early starvation (n=124) and exposed to exogenous DA after 3 days of early starvation (n=63). Each row indicates the age-normalized behavior of one individual across all developmental stages. The different stages are separated by white lines indicating the middle of the lethargus state. Color bar represents the fraction of time spent roaming in each of the 375 time-bins. **(B)** Development time across L1-L4 larval stages within unstarved, starved and starved with exogenous DA *cat-2* populations. **(C)** Average roaming fraction of unstarved, starved and starved with exogenous DA *cat-2* individuals in each developmental stage. Each point represents a single individual. Red bars represent the population mean. **(D)** Average roaming speed of unstarved, starved and starved with exogenous DA *cat-2* individuals in each developmental stage. Each point represents a single individual. Red bars represent the population mean. Upper bars in (C,D) indicate statistical significance (Wilcoxon rank-sum test, FDR corrected) of the difference between unstarved and starved populations in each developmental stage (-log(P-value), indicated are P-values<0.01).

**Figure S5.**
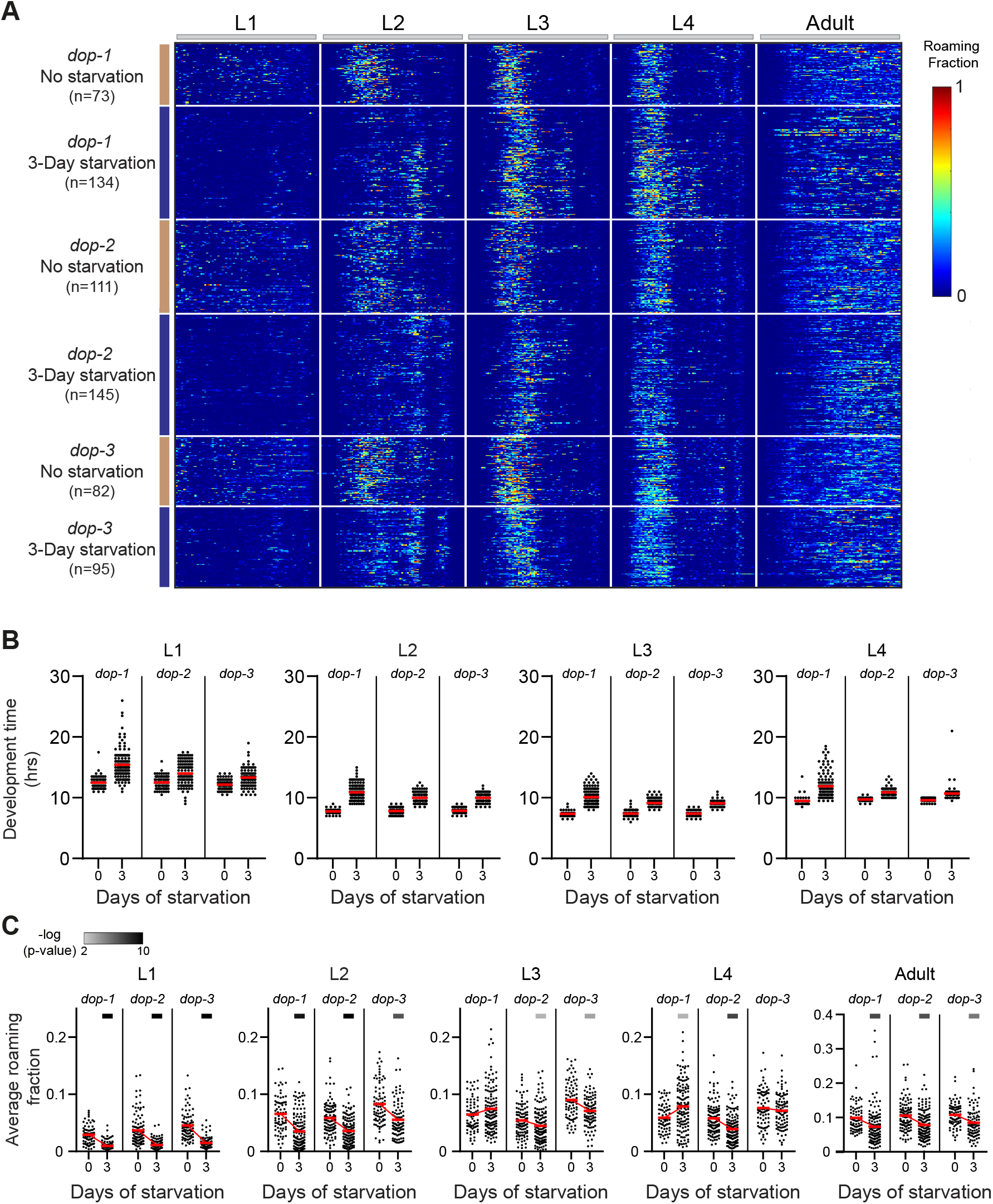
Development-time and roaming quantification in starved and unstarved dopamine receptors mutants. Related to Figure 4. **(A)** Roaming and dwelling behavior of single dopamine receptor mutants without early starvation (*dop-1* n=73; *dop-2* n=111; *dop-3* n= 82) and following 3 days of early starvation (*dop-1* n=134; *dop-2* n=145; *dop-3* n= 95). Each row indicates the age-normalized behavior of one individual across all developmental stages. The different stages are separated by white lines indicating the middle of the lethargus state. Color bar represents the fraction of time spent roaming in each of the 375 time-bins. **(B)** Development time across L1-L4 larval stages within starved and unstarved dopamine receptors mutants. **(C)** Average roaming fraction of starved and unstarved dopamine receptors mutant individuals in each developmental stage. Each point represents a single individual. Red bars represent population mean. Upper bars indicate statistical significance (Wilcoxon rank-sum test, FDR corrected) of the difference between unstarved and starved populations in each developmental stage (-log(P-value), indicated are P-values<0.01).

**Figure S6.**
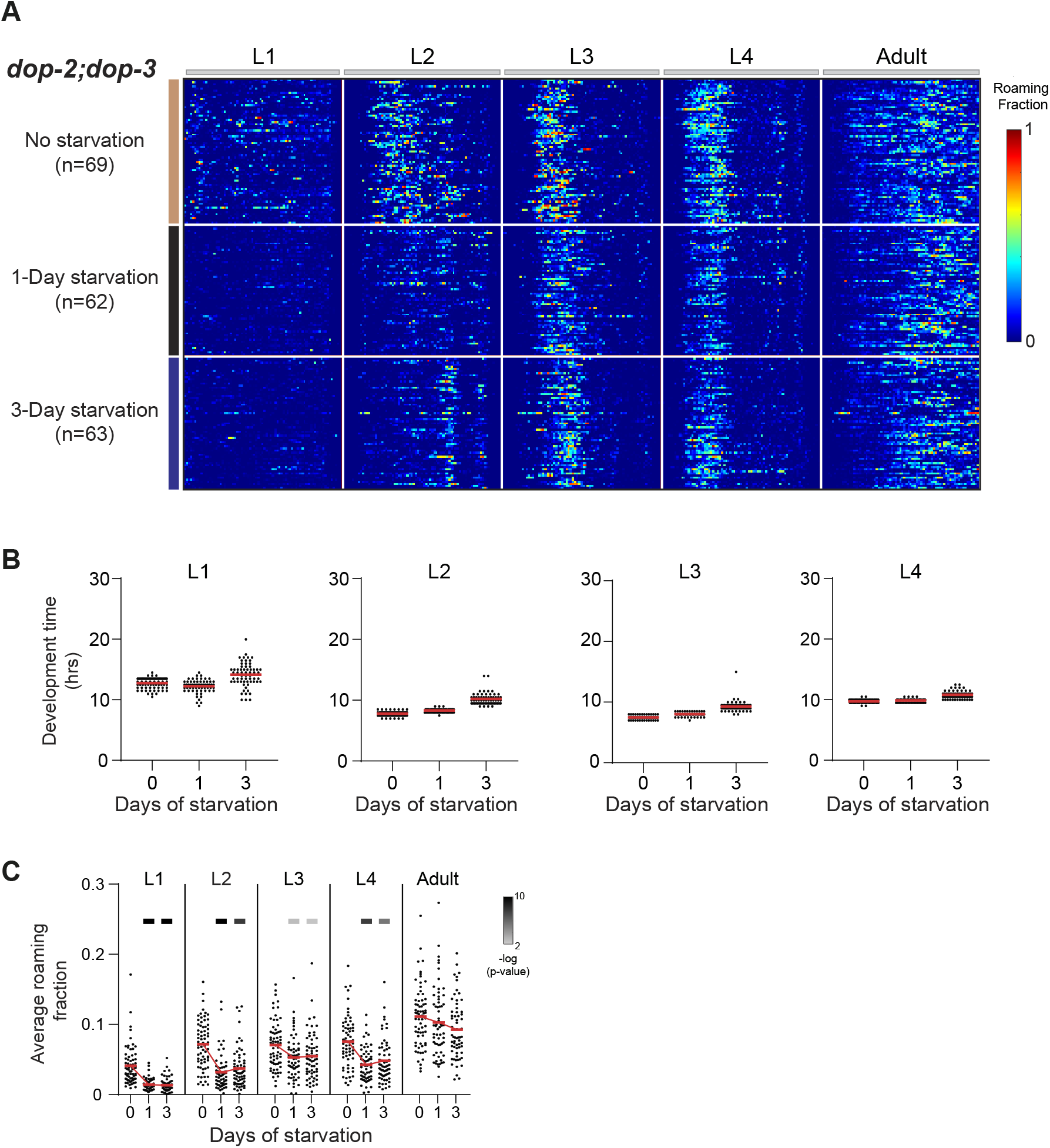
development-time and roaming quantification in starved and unstarved dopamine receptors double mutants. Related to Figure 4. **(A)** Roaming and dwelling behavior of *dop-2;dop-3* animals without early starvation (n=69) and following 1 day (n=62) and 3 days (n=63) of early starvation. Each row indicates the age-normalized behavior of one individual across all developmental stages. The different stages are separated by white lines indicating the middle of the lethargus state. Color bar represents the fraction of time spent roaming in each of the 375 time-bins. **(B)** Development time across L1-L4 larval stages within starved and unstarved *dop-2;dop-3* populations. **(C)** Average roaming fraction of starved and unstarved *dop-2;dop-3* mutant individuals in each developmental stage. Each point represents a single individual. Red bars represent the population mean. Upper bars indicate statistical significance (Wilcoxon rank-sum test, FDR corrected) of the difference between unstarved and starved populations in each developmental stage (-log(P-value), indicated are P-values<0.01).

**Figure S7.**
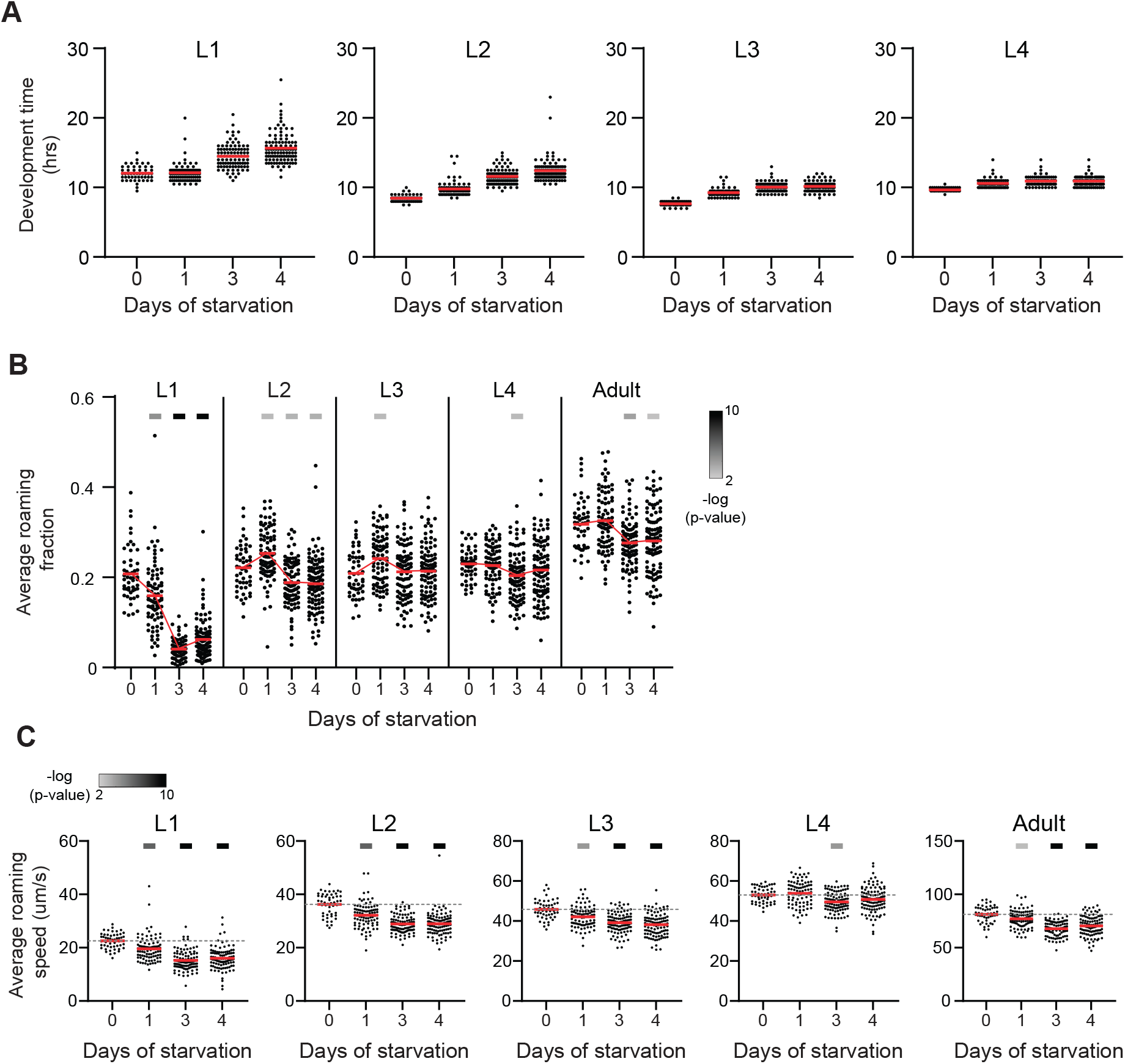
Development-time and roaming quantification in starved and unstarved *tph-1* individuals. Related to Figure 5. **(A)** Development time across L1-L4 larval stages within starved and unstarved *tph-1* populations (no starvation n=51; 1-day starvation n=87; 3-day starvation n=96; 4-day starvation n=104). **(B)** Average roaming fraction of starved and unstarved *tph-1* individuals in each developmental stage. Each point represents a single individual. Red bars represent the population mean. **(C)** Average roaming speed of starved and unstarved *tph-1* individuals in each developmental stage. Each point represents an animal. Red bars represent population mean. Upper bars in (B,C) indicate statistical significance (Wilcoxon rank-sum test, FDR corrected) of the difference between unstarved and starved populations in each developmental stage (-log(P-value), indicated are P-values<0.01).

**Figure S8.**
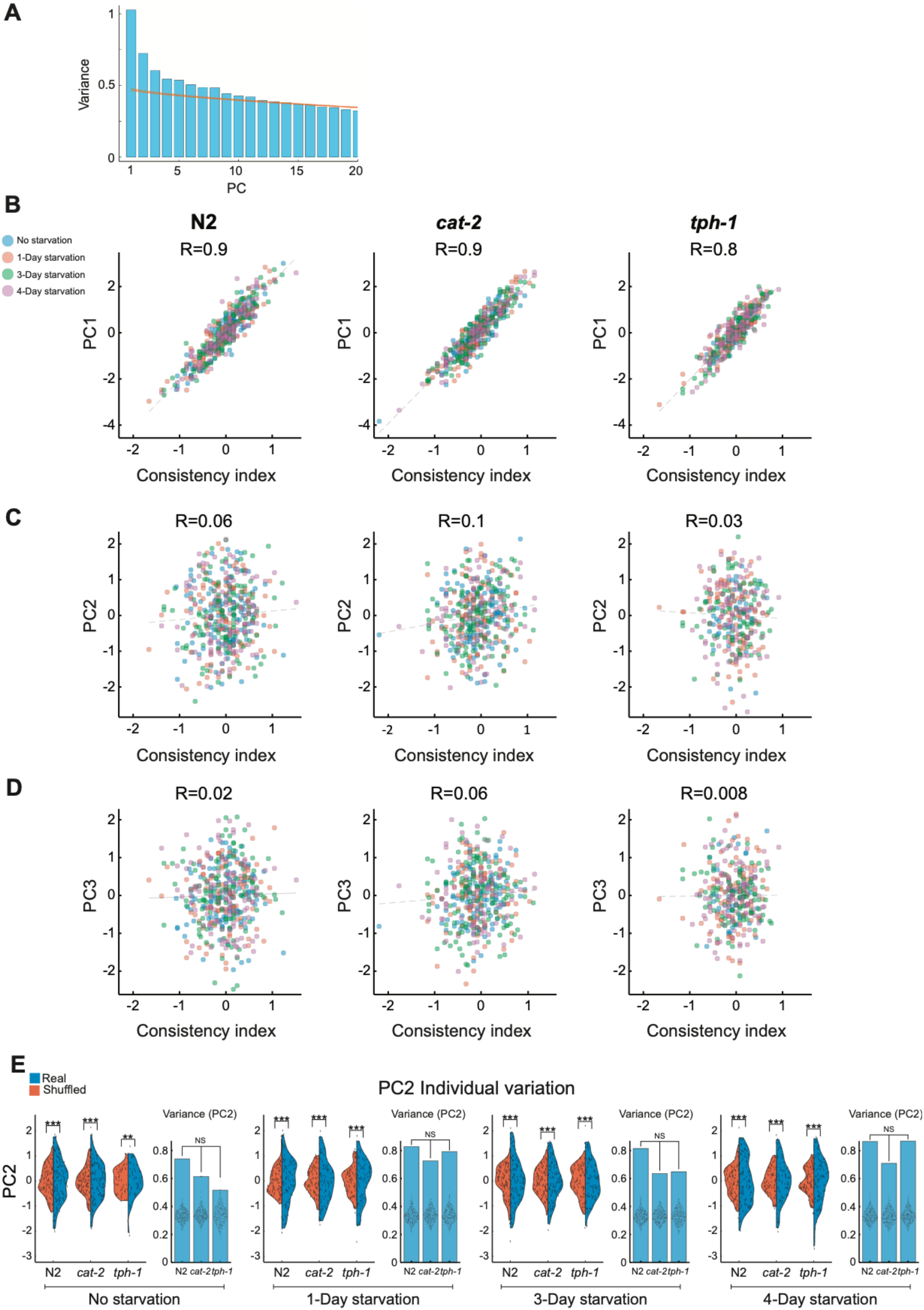
PCA and behavioral consistency analyses in neuromodulatory mutants. Related to Figure 6. **(A)** Variance explained by each of the first 20 PCs following PCA (blue bars) of the pooled wild-type, *tph-1* and *cat-2* individuals rank dataset, compared to the variance explained by the first 20 PCs of a shuffled dataset (500 repetitions, orange lines). **(B-D)** Correlation between behavioral consistency indices and PC1 (B), PC2 (C) and PC3 (D) individual scores within stressed and unstressed wild-type, *tph-1* and *cat-2* populations. Each dot is a single individual, colored by strain and starvation condition. Dotted line is linear least-squares regression with intercept. Pearson correlation coefficient R between behavioral consistency and each PC is noted above the corresponding subplot. **(E)** Distributions of PC2 individual scores (blue) within starved and unstarved wild-type and mutant populations, compared to shuffled dataset (orange). Bar plots represent inter-individual variation in PC2 individual scores. Each dot within bars represents PC2 variation in individual scores within a shuffled dataset (500 repetitions). P-values were calculated using bootstrapping (see Methods). ** P<0.01, ***P<0.001.

## References

Ahamed, T., Costa, A.C., and Stephens, G.J. (2021). Capturing the continuous complexity of behaviour in Caenorhabditis elegans. Nat. Phys. 17, 275–283. https://doi.org/10.1038/s41567-020-01036-8.

Anderson, D.J., and Perona, P. (2014). Toward a Science of Computational Ethology. Neuron 84, 18–31. https://doi.org/10.1016/j.neuron.2014.09.005.

Arous, J.B., Laffont, S., and Chatenay, D. (2009). Molecular and Sensory Basis of a Food Related Two-State Behavior in C. elegans. PLOS ONE 4, e7584. https://doi.org/10.1371/journal.pone.0007584.

Aton, S.J., Colwell, C.S., Harmar, A.J., Waschek, J., and Herzog, E.D. (2005). Vasoactive intestinal polypeptide mediates circadian rhythmicity and synchrony in mammalian clock neurons. Nat Neurosci 8, 476–483. https://doi.org/10.1038/nn1419.

Bargmann, C.I. (2012). Beyond the connectome: How neuromodulators shape neural circuits. Bioessays 34, 458–465. https://doi.org/10.1002/bies.201100185.

Baugh, L.R. (2013). To Grow or Not to Grow: Nutritional Control of Development During *Caenorhabditis elegans* L1 Arrest. Genetics 194, 539–555. https://doi.org/10.1534/genetics.113.150847.

Bierbach, D., Laskowski, K.L., and Wolf, M. (2017). Behavioural individuality in clonal fish arises despite near-identical rearing conditions. Nat Commun 8, 15361. https://doi.org/10.1038/ncomms15361.

Brittin, C.A., Cook, S.J., Hall, D.H., Emmons, S.W., and Cohen, N. (2021). A multi-scale brain map derived from whole-brain volumetric reconstructions. Nature 591, 105–110. https://doi.org/10.1038/s41586-021-03284-x.

Brown, A.E.X., and de Bivort, B. (2018). Ethology as a physical science. Nature Phys 14, 653–657. https://doi.org/10.1038/s41567-018-0093-0.

Buchanan, S.M., Kain, J.S., and de Bivort, B.L. (2015). Neuronal control of locomotor handedness in Drosophila. Proc. Natl. Acad. Sci. U.S.A. 112, 6700–6705. https://doi.org/10.1073/pnas.1500804112.

Casanueva, M.O., Burga, A., and Lehner, B. (2012). Fitness Trade-Offs and Environmentally Induced Mutation Buffering in Isogenic *C. elegans*. Science 335, 82–85. https://doi.org/10.1126/science.1213491.

Cassada, R.C., and Russell, R.L. (1975). The dauerlarva, a post-embryonic developmental variant of the nematode Caenorhabditis elegans. Developmental Biology 46, 326–342. https://doi.org/10.1016/0012-1606(75)90109-8.

Cermak, N., Yu, S.K., Clark, R., Huang, Y.-C., Baskoylu, S.N., and Flavell, S.W. (2020). Whole-organism behavioral profiling reveals a role for dopamine in state-dependent motor program coupling in C. elegans. ELife 9, e57093. https://doi.org/10.7554/eLife.57093.

Chase, D.L., Pepper, J.S., and Koelle, M.R. (2004). Mechanism of extrasynaptic dopamine signaling in Caenorhabditis elegans. Nat Neurosci 7, 1096–1103. https://doi.org/10.1038/nn1316.

Churgin, M.A., Lavrentovich, D., Smith, M.A., Gao, R., Boyden, E., and de Bivort, B. (2021). Neural correlates of individual odor preference in *Drosophila* (Neuroscience).

Cooper, W.S., and Kaplan, R.H. (1982). Adaptive “coin-flipping”: a decision-theoretic examination of natural selection for random individual variation. J Theor Biol 94, 135–151. https://doi.org/10.1016/0022-5193(82)90336-8.

Flavell, S.W., Pokala, N., Macosko, E.Z., Albrecht, D.R., Larsch, J., and Bargmann, (2013). Serotonin and the Neuropeptide PDF Initiate and Extend Opposing Behavioral States in C. elegans. Cell 154, 1023–1035. https://doi.org/10.1016/j.cell.2013.08.001.

Freund, J., Brandmaier, A.M., Lewejohann, L., Kirste, I., Kritzler, M., Kruger, A., Sachser, N., Lindenberger, U., and Kempermann, G. (2013). Emergence of Individuality in Genetically Identical Mice. Science 340, 756–759. https://doi.org/10.1126/science.1235294.

Fujiwara, M., Sengupta, P., and McIntire, S.L. (2002). Regulation of Body Size and Behavioral State of C. elegans by Sensory Perception and the EGL-4 cGMP-Dependent Protein Kinase. Neuron 36, 1091–1102. https://doi.org/10.1016/S0896-6273(02)01093-0.

Greenwald, I.S., and Horvitz, H.R. (1982). DOMINANT SUPPRESSORS OF A MUSCLE MUTANT DEFINE AN ESSENTIAL GENE OF *CAENORHABDITIS ELEGANS*. Genetics 101, 211–225. https://doi.org/10.1093/genetics/101.2.211.

Harris-Warrick, R.M., and Marder, E. (1991). Modulation of Neural Networks for Behavior. Annu. Rev. Neurosci. 14, 39–57. https://doi.org/10.1146/annurev.ne.14.030191.000351.

Honegger, K., and de Bivort, B. (2018). Stochasticity, individuality and behavior. Current Biology 28, R8–R12. https://doi.org/10.1016/j.cub.2017.11.058.

Honegger, K.S., Smith, M.A.-Y., Churgin, M.A., Turner, G.C., and de Bivort, B.L. (2020). Idiosyncratic neural coding and neuromodulation of olfactory individuality in *Drosophila*. Proc Natl Acad Sci USA 117, 23292–23297. https://doi.org/10.1073/pnas.1901623116.

Horn, G. (1998). Visual imprinting and the neural mechanisms of recognition memory. Trends in Neurosciences 21, 300–305. https://doi.org/10.1016/S0166-2236(97)01219-8.

Immelmann, K. (1975). Ecological significance of imprinting and early learning. Annu. Rev. Ecol. Syst. 6, 15–37. .

Jin, X., Pokala, N., and Bargmann, C.I. (2016). Distinct Circuits for the Formation and Retrieval of an Imprinted Olfactory Memory. Cell 164, 632–643. https://doi.org/10.1016/j.cell.2016.01.007.

Johnson, T.E., Mitchell, D.H., Kline, S., Kemal, R., and Foy, J. (1984). Arresting development arrests aging in the nematode Caenorhabditis elegans. Mechanisms of Ageing and Development 28, 23–40. https://doi.org/10.1016/0047-6374(84)90150-7.

Kain, J.S., Stokes, C., and de Bivort, B.L. (2012). Phototactic personality in fruit flies and its suppression by serotonin and white. Proceedings of the National Academy of Sciences 109, 19834–19839. https://doi.org/10.1073/pnas.1211988109.

Kennedy, A., Asahina, K., Hoopfer, E., Inagaki, H., Jung, Y., Lee, H., Remedios, R., and Anderson, D.J. (2014). Internal States and Behavioral Decision-Making: Toward an Integration of Emotion and Cognition. Cold Spring Harb Symp Quant Biol 79, 199–210. https://doi.org/10.1101/sqb.2014.79.024984.

Kimmel, C.B., Patterson, J., and Kimmel, R.O. (1974). The development and behavioral characteristics of the startle response in the zebra fish. Dev. Psychobiol. 7, 47–60. https://doi.org/10.1002/dev.420070109.

Korosi, A., Naninck, E.F.G., Oomen, C.A., Schouten, M., Krugers, H., Fitzsimons, C., and Lucassen, P.J. (2012). Early-life stress mediated modulation of adult neurogenesis and behavior. Behavioural Brain Research 227, 400–409. https://doi.org/10.1016/j.bbr.2011.07.037.

Linneweber, G.A., Andriatsilavo, M., Dutta, S.B., Bengochea, M., Hellbruegge, L., Liu, G., Ejsmont, R.K., Straw, A.D., Wernet, M., Hiesinger, P.R., et al. (2020). A neurodevelopmental origin of behavioral individuality in the Drosophila visual system. Science 367, 1112–1119. https://doi.org/10.1126/science.aaw7182.

Lints, R., and Emmons, S.W. (1999). Patterning of dopaminergic neurotransmitter identity among Caenorhabditis elegans ray sensory neurons by a TGFbeta family signaling pathway and a Hox gene. Development 126, 5819–5831. https://doi.org/10.1242/dev.126.24.5819.

Lorenz, K. (1935). Der Kumpan in der Umwelt des Vogels: Der Artgenosse als auslösendes Moment sozialer Verhaltungsweisen. J. Ornithol 83, 137–213. https://doi.org/10.1007/BF01905355.

Marder, E. (2012). Neuromodulation of Neuronal Circuits: Back to the Future. Neuron 76, 1–11. https://doi.org/10.1016/j.neuron.2012.09.010.

Marder, E., Kedia, S., and Morozova, E.O. (2022). New insights from small rhythmic circuits. Current Opinion in Neurobiology 76, 102610. https://doi.org/10.1016/j.conb.2022.102610.

Marella, S., Mann, K., and Scott, K. (2012). Dopaminergic Modulation of Sucrose Acceptance Behavior in Drosophila. Neuron 73, 941–950. https://doi.org/10.1016/j.neuron.2011.12.032.

Nakamori, T., Maekawa, F., Sato, K., Tanaka, K., and Ohki-Hamazaki, H. (2013). Neural basis of imprinting behavior in chicks. Develop. Growth Differ. 55, 198–206. https://doi.org/10.1111/dgd.12028.

Nevitt, G.A., Dittman, A.H., Quinn, T.P., and Moody, W.J. (1994). Evidence for a peripheral olfactory memory in imprinted salmon. Proceedings of the National Academy of Sciences 91, 4288–4292. https://doi.org/10.1073/pnas.91.10.4288.

Nusbaum, M.P., and Blitz, D.M. (2012). Neuropeptide modulation of microcircuits. Current Opinion in Neurobiology 22, 592–601. https://doi.org/10.1016/j.conb.2012.01.003.

Omura, D.T., Clark, D.A., Samuel, A.D.T., and Horvitz, H.R. (2012). Dopamine Signaling Is Essential for Precise Rates of Locomotion by C. elegans. PLoS ONE 7, e38649. https://doi.org/10.1371/journal.pone.0038649.

Pantoja, C., Hoagland, A., Carroll, E.C., Karalis, V., Conner, A., and Isacoff, E.Y. (2016). Neuromodulatory Regulation of Behavioral Individuality in Zebrafish. Neuron 91, 587–601. https://doi.org/10.1016/j.neuron.2016.06.016.

Park, J.H., and Hall, J.C. (1998). Isolation and Chronobiological Analysis of a Neuropeptide Pigment-Dispersing Factor Gene in *Drosophila melanogaster*. J Biol Rhythms 13, 219–228. https://doi.org/10.1177/074873098129000066.

Pattwell, S.S., Duhoux, S., Hartley, C.A., Johnson, D.C., Jing, D., Elliott, M.D., Ruberry, E.J., Powers, A., Mehta, N., Yang, R.R., et al. (2012). Altered fear learning across development in both mouse and human. Proceedings of the National Academy of Sciences 109, 16318–16323. https://doi.org/10.1073/pnas.1206834109.

Pradhan, S., Quilez, S., Homer, K., and Hendricks, M. (2019). Environmental Programming of Adult Foraging Behavior in C. elegans. Current Biology 29, 2867–2879.e4. https://doi.org/10.1016/j.cub.2019.07.045.

Rehm, K.J., Deeg, K.E., and Marder, E. (2008). Developmental Regulation of Neuromodulator Function in the Stomatogastric Ganglion of the Lobster, Homarus americanus. Journal of Neuroscience 28, 9828–9839. https://doi.org/10.1523/JNEUROSCI.2328-08.2008.

Remy, J.-J., and Hobert, O. (2005). An Interneuronal Chemoreceptor Required for Olfactory Imprinting in *C. elegans*. Science 309, 787–790. https://doi.org/10.1126/science.1114209.

Sanyal, S., Wintle, R.F., Kindt, K.S., Nuttley, W.M., Arvan, R., Fitzmaurice, P., Bigras, E., Merz, D.C., Hébert, T.E., van der Kooy, D., et al. (2004). Dopamine modulates the plasticity of mechanosensory responses in Caenorhabditis elegans. EMBO J 23, 473–482. https://doi.org/10.1038/sj.emboj.7600057.

Sasaki-Adams, D. (2001). Serotonin-Dopamine Interactions in the Control of Conditioned Reinforcement and Motor Behavior. Neuropsychopharmacology 25, 440–452. https://doi.org/10.1016/S0893-133X(01)00240-8.

Sawin, E.R., Ranganathan, R., and Horvitz, H.R. (2000). C. elegans Locomotory Rate Is Modulated by the Environment through a Dopaminergic Pathway and by Experience through a Serotonergic Pathway. Neuron 26, 619–631. https://doi.org/10.1016/S0896-6273(00)81199-X.

Schuett, W., Dall, S.R.X., Baeumer, J., Kloesener, M.H., Nakagawa, S., Beinlich, F., and Eggers, T. (2011). Personality variation in a clonal insect: the pea aphid, Acyrthosiphon pisum. Dev Psychobiol 53, 631–640. https://doi.org/10.1002/dev.20538.

Schwarz, R.F., Branicky, R., Grundy, L.J., Schafer, W.R., and Brown, A.E.X. (2015). Changes in Postural Syntax Characterize Sensory Modulation and Natural Variation of C. elegans Locomotion. PLoS Comput Biol 11, e1004322. https://doi.org/10.1371/journal.pcbi.1004322.

Sisk, C.L., and Foster, D.L. (2004). The neural basis of puberty and adolescence. Nat Neurosci 7, 1040–1047. https://doi.org/10.1038/nn1326.

Sokolowski, M.B., Kent, C., and Wong, J. (1984). Drosophila larval foraging behaviour: Developmental stages. Animal Behaviour 32, 645–651. https://doi.org/10.1016/S0003-3472(84)80139-6.

Stern, S., Kirst, C., and Bargmann, C.I. (2017). Neuromodulatory Control of Long-Term Behavioral Patterns and Individuality across Development. Cell 171, 1649–1662.e10. https://doi.org/10.1016/j.cell.2017.10.041.

Sugiura, M., Fuke, S., Suo, S., Sasagawa, N., Van Tol, H.H.M., and Ishiura, S. (2005). Characterization of a novel D2-like dopamine receptor with a truncated splice variant and a D1-like dopamine receptor unique to invertebrates from Caenorhabditis elegans: Dopamine receptors from C. elegans. Journal of Neurochemistry 94, 1146–1157. https://doi.org/10.1111/j.1471-4159.2005.03268.x.

Sulston, J., Dew, M., and Brenner, S. (1975). Dopaminergic neurons in the nematodeCaenorhabditis elegans. J. Comp. Neurol. 163, 215–226. https://doi.org/10.1002/cne.901630207.

Suo, S., Sasagawa, N., and Ishiura, S. (2003). Cloning and characterization of a Caenorhabditis elegans D2-like dopamine receptor: C. elegans D2-like dopamine receptor. Journal of Neurochemistry 86, 869–878. https://doi.org/10.1046/j.1471-4159.2003.01896.x.

Suo, S., Culotti, J.G., and Van Tol, H.H.M. (2009). Dopamine counteracts octopamine signalling in a neural circuit mediating food response in C. elegans. EMBO J 28, 2437–2448. https://doi.org/10.1038/emboj.2009.194.

Taghert, P.H., and Nitabach, M.N. (2012). Peptide Neuromodulation in Invertebrate Model Systems. Neuron 76, 82–97. https://doi.org/10.1016/j.neuron.2012.08.035.

Tang, L.S., Taylor, A.L., Rinberg, A., and Marder, E. (2012). Robustness of a Rhythmic Circuit to Short- and Long-Term Temperature Changes. Journal of Neuroscience 32, 10075–10085. https://doi.org/10.1523/JNEUROSCI.1443-12.2012.

Truman, J.W. (2005). Hormonal Control of Insect Ecdysis: Endocrine Cascades for Coordinating Behavior with Physiology. In Vitamins & Hormones, (Elsevier), pp. 1–30.

Tsalik, E.L., Niacaris, T., Wenick, A.S., Pau, K., Avery, L., and Hobert, O. (2003). LIM homeobox gene-dependent expression of biogenic amine receptors in restricted regions of the C. elegans nervous system. Developmental Biology 263, 81–102. https://doi.org/10.1016/S0012-1606(03)00447-0.

Wigglesworth, V.B. (1936). Memoirs: The function of the Corpus Allatum in the Growth and Reproduction of Rhodnius Prolixus (Hemiptera). Journal of Cell Science s2-79, 91–121. https://doi.org/10.1242/jcs.s2-79.313.91.

Wilson, D.A., and Sullivan, R.M. (1994). Neurobiology of associative learning in the neonate: Early olfactory learning. Behavioral and Neural Biology 61, 1–18. https://doi.org/10.1016/S0163-1047(05)80039-1.

Witvliet, D., Mulcahy, B., Mitchell, J.K., Meirovitch, Y., Berger, D.R., Wu, Y., Liu, Y., Koh, W.X., Parvathala, R., Holmyard, D., et al. (2021). Connectomes across development reveal principles of brain maturation. Nature 596, 257–261. https://doi.org/10.1038/s41586-021-03778-8.

Zhang, Y., Lu, H., and Bargmann, C.I. (2005). Pathogenic bacteria induce aversive olfactory learning in Caenorhabditis elegans. Nature 438, 179–184. https://doi.org/10.1038/nature04216.

